# A robust and user-agnostic step-emulsion platform for scalable microgel fabrication

**DOI:** 10.64898/2026.05.05.722106

**Authors:** Durante Pioche-Lee, Shiyu Yang, Xiaoqian Wang, Yoke Qi Ho, Wahidur Rahman, Armen C. Vartanian, Despina I. Pavlidis, Irene W. Zhang, Julia E. Vallier, Emily McCorkle, Andrew Schaefer, Andrew J. Putnam, Ariella Shikanov, Cole A. DeForest, Sasha Cai Lesher-Pérez

## Abstract

Over the past decade, the integration of microgel-based granular hydrogels in biomedical technologies has experienced substantial growth due to the numerous benefits microgels offer. However, the inability to easily adopt uniform microgel fabrication workflows at scale constitutes a major bottleneck, or in some cases, a barrier-to-entry that stunts further growth of the field. The gold-standard technique for emulsion-based microgel production is through microfluidic droplet-generating devices that produce liquid gel precursor droplets that gel post-production. However, traditional microfluidic workflows often require multiple independent flows and controlled pressure sources, along with a steep learning curve in using microfluidics to achieve uniform droplet sizes reproducibly and repeatedly. This difficulty in adopting microgel fabrication is further compounded by low throughput and the extensive flow rate calibration required when switching to new formulations (e.g., material type, droplet size). In this work, we present a step-emulsion system that bridges the gap by providing a robust and simple setup. We experimentally characterize and evaluate how flow and outlet channel dimension contribute to the generation of uniform droplet populations at specific sizes. With our large dataset consisting of various outlet channel dimensions, we evaluated outlet channel geometrical impacts (height, width, cross-sectional area, aspect-ratio, etc.) on gel precursor droplet size and generation throughput. We demonstrate robust, highly compatible, and repeatably uniform droplet generation from various gel precursor polymer backbones, users with varying microfluidics experience, and a wide viscosity range, including alginate solutions with 650 times the viscosity of water. Furthermore, we confirmed consistent gel precursor droplet generation outcomes driven by a constant flow source (syringe pump) and by direct manual injection as a simple and highly adoptable option for the generation of gel precursor droplets. This platform is ideal for researchers seeking rapid and easy microgel fabrication, regardless of microfluidics experience.

## 1. Introduction

Microgels, and their composite structures of granular hydrogels, have gained popularity due to the combined benefit of being both injectable and achieving microscale porosity when packed together. The inherent modularity of microgels as building blocks enables tuning of the constituent microgel characteristics (e.g., stiffness, degradability, surface modification), while decoupling the porosity of the granular hydrogels. This new class of biomaterials has multifold applications, most notably in wound healing and regenerative medicine due to improved integration into host tissues as compared to solid, bulk hydrogels.^[1–4]^ More recently, these microgel-based materials have been used as bioinks^[5,6]^ and as suspension baths for bioprinting^[7–9]^ to produce in vitro cell scaffolds, tissue engineered constructs, and as implantable units.

However, a key challenge for both industrial and academic research applications of microgels is achieving reliable, uniform production at scale, which ensures reproducibility among the intended downstream applications.^[10–12]^ While there are multiple methods to generate microgels, likely the most ubiquitous is producing emulsions of gel precursor droplets within an oil solution, wherein the droplets can subsequently be polymerized into microgels through spontaneous and/or exogenously triggered crosslinking chemistries. The current gold standard for producing monodisperse droplet populations, and consequently uniform microgels, is using microfluidic droplet generators. These systems may be widespread in the literature, however, obtaining sufficient microfluidic knowledge to be well-versed in the know-how and tricks, as well as needing the appropriate equipment to run these systems can present an activation barrier for non-microfluidicists in order to achieve easy adoption and immediate, successful droplet generation^[11]^ and microgel production. This barrier is further enhanced when the needed volume reaches the milliliter range, as many of the gold standard droplet generators (e.g., flow-focusing, co-flow, cross-flow devices) used for microgel production are low throughput^[13–17]^, resulting in a bottleneck to produce the desired droplet populations at scale.

Higher-throughput droplet generator systems, such as step-emulsion devices, have been deployed by various groups for microgel production.^[15,18–20]^ Step emulsification in these devices produces droplets by generating a Laplace pressure difference as the dispersed phase exits the confined outlet channel to an expanded reservoir, resulting in a neck and bulb until the bulb detaches (as a droplet) when the neck reaches a critical thinning width.^[14,15,17,20–22]^ Step-emulsion device designs offer an attractive alternative, as they can increase droplet generation by orders of magnitude over traditional microfluidic droplet generators through the massive parallelization of the droplet generation channels.^[17]^ In configurations that employ a clearing, continuous sheath phase^[23–25]^ to move any formed droplets away from the production site (i.e., channel outlet), additional equipment is needed to independently control both the continuous and dispersed phase flow rates, increasing system complexity.

While there are numerous benefits of these systems, many step-emulsion droplet generators are structured serially, such that a defect or disruption at any point in the device can propagate downstream and lead to non-uniform droplet production, increased polydispersity, or device failure. Moreover, serial designs that rely on a flowing continuous phase to clear droplets increase the total hydraulic resistance, thereby increasing operating pressure and requiring further flow rate tuning to mitigate backpressure-induced increases in droplet polydispersity. Additionally, changes in gel precursor properties (e.g., viscosity) often require additional flow rate optimization of both the dripping and continuous phases for successful droplet generation.

Furthermore, step-emulsion devices in literature are primarily used to produce a limited range of droplet sizes, often utilizing only a single gel precursor formulation per study, and require a minimum of two syringe pumps to independently control both the aqueous droplet-generation flow and the oil-clearing flow rates. While well-described methods exist to facilitate easier adoption of these systems^[26]^, the limited guidance in literature for redesigning these systems to adjust sizes or accommodate solutions of distinct viscosities creates a substantial barrier for simple adoption and modification for non-microfluidic specialists. We are acutely aware of these barriers in the microgel and granular biomaterial’s community, though we expect the barriers to be broadly experienced across multiple fields (e.g., pharmaceuticals, food, agriculture, cosmetics, etc.,) that aim to leverage droplet generation.

In this work, we present a modular device design for droplet generation of gel precursor solutions for microgel production. To achieve an easy-to-use platform and thereby democratize droplet-based research, we build on traditional step-emulsion device configurations to produce a new device, which is submerged in a reservoir of fluorinated oil of higher density relative to the aqueous gel precursor and only requires inflow of the gel precursor (**Figure 1A**). The gel precursor flows out of the device laterally and buoyancy clears the formed droplets away from the droplet generation channel exit, floating to the top surface of the reservoir where subsequent gelation (triggered or spontaneous) and collection of the microgels can occur (Figure 1B-C). Images of the device during droplet generation can be seen in **Figure S1** and videos of droplet generation are included as additional files. Critically, this device can be used with multiple gelation chemistries utilizing various common biomaterial backbones (Figure 1D), including alginate, hyaluronic acid (HA), gelatin, and poly(ethylene glycol) (PEG), and produces similar droplet sizes, as monodispersed populations across a range of flow rates (Figure 1E-F, and **Figure S2**).

**Figure 1.**
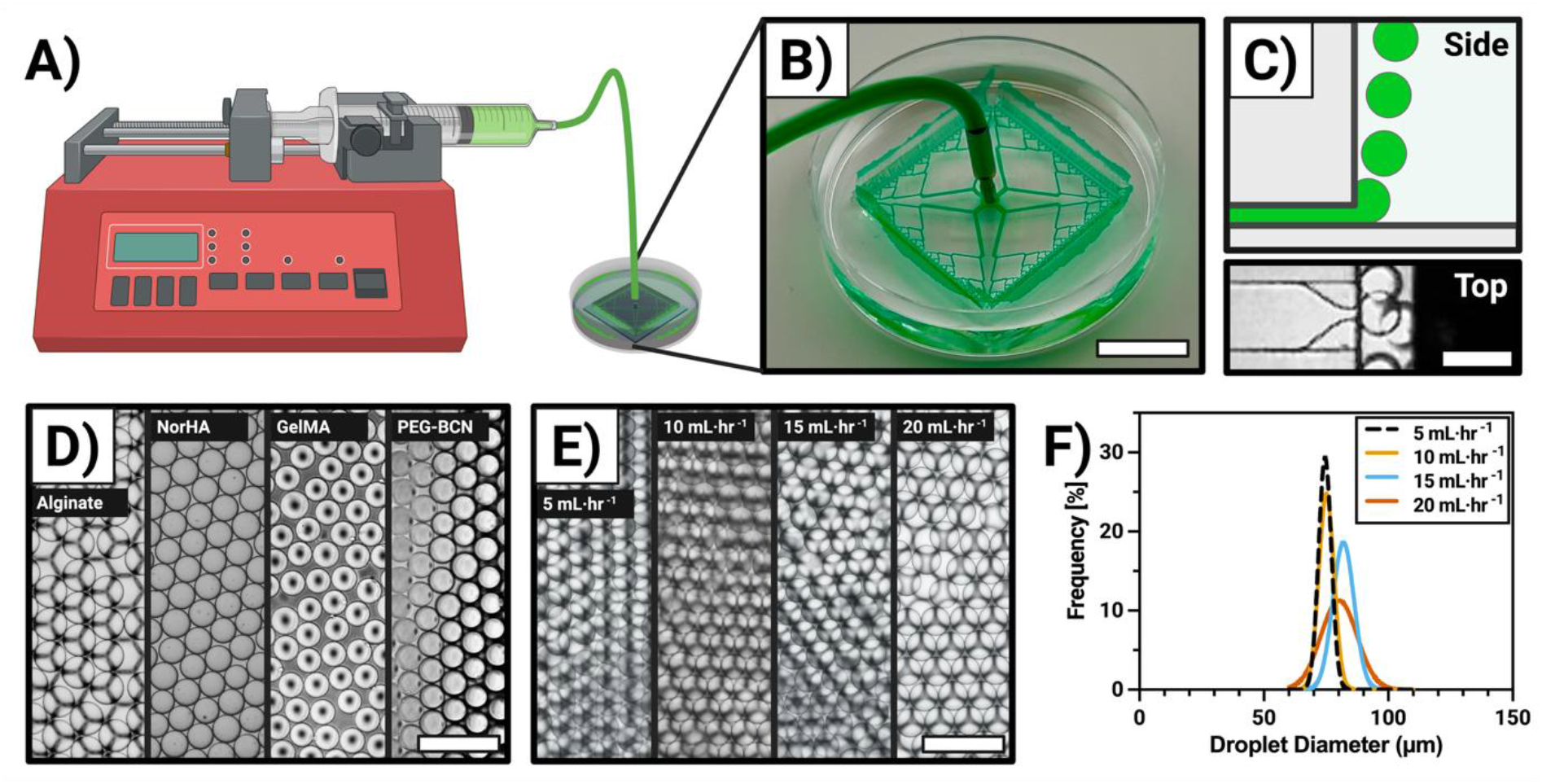
A) Schematic overview of droplet generation using the step-emulsion buoyancy-clearing device for microgel. B) Parallelized outlet channels generating droplets enable increased throughput (schematic shows 512-outlet channel device with 30 µm height and 150 µm width producing droplets of water with food dye droplets for visualization); scale bar is 15 mm. C) Representative side-view of droplet generation and top-view image of PEG-NB hydrogel precursor droplet formation from a 512-outlet channel device with outlet channel dimensions of 20 µm height and 120 µm width; scale bar is 120 µm. D) The droplet generation system was tested with common biomaterial backbones including alginate, hyaluronic acid-norbornene, gelatin-methacrylate, and poly(ethylene glycol) (PEG)-BCN, and is largely independent of the material-type. Images of alginate, hyaluronic acid-norbornene, and gelatin-methacrylate are shown as droplets prior to gelation. PEG-BCN droplets immediately gel post-production, and therefore PEG-BCN images show microgels. Images are from a 512-outlet channel device with outlet channel dimensions of 20 µm height and 120 µm width. All devices were infused at 10 mL hr^−1^; scale bar is 250 µm. E & F) The system is also largely independent of flow, producing similar PEG-norbornene (PEG-NB) hydrogel precursor droplet populations at various flow rates, shown by Gaussian distributions of the collected droplets from a 512-outlet channel device with outlet channel dimensions of 20 µm height and 120 µm width. Images of PEG-norbornene (PEG-NB) show gel precursor droplets (prior to gelation) and were generated by infusing the device with 10 wt% PEG-NB hydrogel precursor solution. Scatter dot plots can be found in Figure S2; scale bar is 250 µm.

To assess the robustness of our device design, we fabricated a library of droplet generators capable of producing monodisperse droplets across a range of sizes defined by channel geometry. We confirmed aspect ratio (i.e., channel height over width) impacted both droplet size and range of flow rates for monodispersed droplet production, as has been previously reported in other step-emulsion systems^[18]^. To the best of our knowledge, this work is the first to experimentally evaluate multiple aspect ratio characterization in a lateral step-emulsion system. We determined that increasing aspect ratio while controlling for cross-sectional area increased the maximum flow velocities that could be run prior to transitioning from monodispersed to polydispersed populations. Furthermore, to provide insight to the broader community on what channel characteristics most impacted droplet size, we evaluated a range of step-emulsion geometries and generated droplets ranging from approximately 50 µm to 600 µm. Our analysis revealed that droplet size correlated to the smallest dimension of the channel (either width or height), rather than height alone. Unlike previously reported step-emulsion systems that leverage buoyancy clearance and generate droplets in the vertical direction^[15,20,27,28]^, our lateral droplet generation facilitates a radial design around the aqueous inflow, parallelizing the droplet generators and distributing pressure and flow velocity uniformly across the device. Based on our radial design and the parallelized 512-droplet generation channels, we produced monodisperse microgels at 10 mL hr^−1^ across various common gel precursor solutions using different backbones and gelation chemistries, i.e., PEG-norbornene, PEG-diacrylate, PEG-tetrabicyclononyne, PEG-maleimide, gelatin-methacrylate, gelatin-norbornene, HA-norbornene, and alginate.

We further demonstrated that due to the robustness across a range of flow rates, this device could be operated either via syringe pump or direct manual injection of the fluid, producing monodispersed droplets of similar sizes in both modes. As further evidence of the robustness and simplicity of the system, multiple users with varying microfluidics experience used the devices, achieving successful, similarly sized, monodispersed droplet generation using both syringe pump-driven and manual injection methods. We additionally evaluated the role that higher viscosity solutions could play within droplet generation and demonstrated the ability to produce monodispersed droplets with solution viscosities approximately 650 times greater than water, though we had to adjust flow rates to fall below the critical capillary number^[19,20]^ to stay within a dripping regime. Fundamentally, our work presents a simple, calibration-free, plug-and-play microfluidic platform for high-throughput monodispersed gel precursor droplet production as well as a workflow for easy adoption by researchers of all experience levels to expedite the production of monodispersed microgels at large scales.

## 2. Results and Discussion

In this work, droplet generation was achieved using either a 32- or 512-outlet channel device design using various synthetic polymers and common biomaterials. However, unless biomaterial composition is explicitly noted, the system was characterized using a gel precursor solution of 10 wt% poly(ethylene glycol) norbornene (PEG-NB, 8-arm, 40 kDa molecular weight) at a 0.4 r-ratio with poly(ethylene) glycol dithiol with lithium phenyl-2,4,6-trimethylbenzoylphosphinate (LAP, 2.2 mM) as the photoinitiator. It is critical to note that all the data we present is for gel precursor droplets prior to gelation, except in the case of PEG-BCN, which immediately gelled post-droplet production via the spontaneous strain-promoted azide-alkyne cycloaddition with a PEG-diazide. Droplets were specifically assessed, as we identified no difference between pre-gelation droplets and the corresponding post-gelation microgels while in the fluorinated oil. In all cases, hydrogel precursor droplets underwent gelation and were subsequently purified to verify no issues with the droplet fabrication and gelation process with our system and the fluorinated oils. Data on purified microgels is not presented, as during the purification process, microgels can begin to swell. Evaluation of purified microgels would thus confound comparison of the droplet generator, as swelling is an independent characteristic of the different materials (i.e., hydrogel compositions) used.

### 2.1 Flow rate and aspect ratio influence droplet generation

#### 2.1.1 Higher aspect ratio channels maintain droplet generation stability at higher flow velocities

In step-emulsion systems, channel geometry has been identified to play an important role in determining both droplet size and a system’s ability to generate droplets in the dripping regime.^[18,24,29,30]^ When step-emulsion systems are operated in a stable droplet producing flow regime, droplet generation size only depends on outlet channel geometry and is independent of flow rate^[19]^ resulting in monodisperse droplets. Here, we evaluated the impact of increasing flow rates on uniform droplet generation for different aspect ratios – height over width (h/w) – in our step-emulsion device, while keeping similar cross-sectional areas for the channels (Figure 2A). To the best of our knowledge, this has not been extensively evaluated for lateral step-emulsion devices.

**Figure 2.**
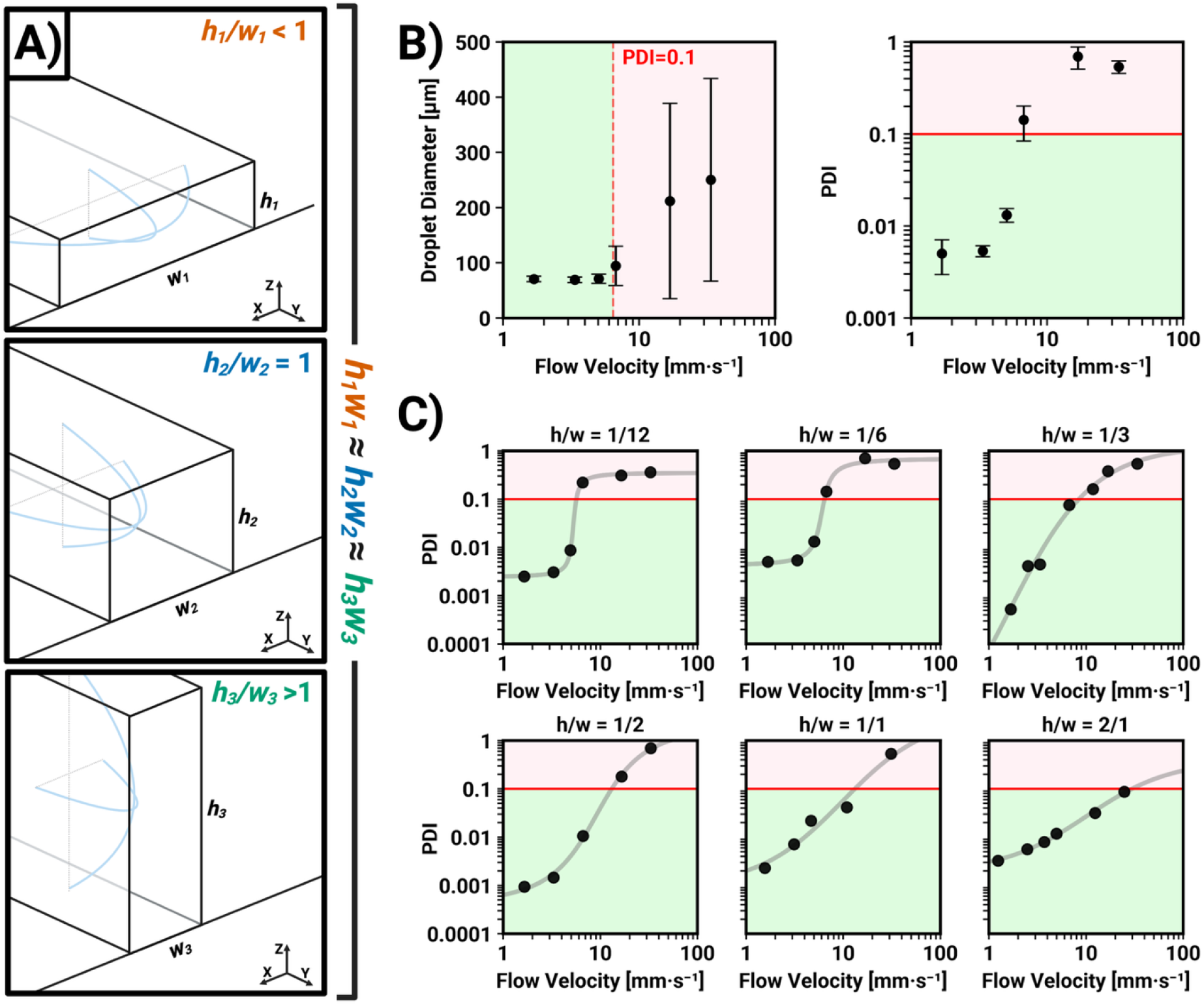
Greater height-to-width (h/w) aspect ratios with constant cross-sectional area maintain stable flow regime at higher flow velocities for our step-emulsion droplet generator. A) Schematic of different h/w aspect ratio geometries with constant cross-sectional area. B) In the stable regime (highlighted in green), flow velocity results in small changes in droplet size and PDI (device geometry h/w = 1/6, 32-outlet channels, dispersed phase is 10 wt% PEG-NB). Increasing above a certain flow velocity (highlighted in red), dispersity (PDI) transitions out of the monodisperse (PDI < 0.1, transition determined by logistical regression) droplet producing regime. C) Increasing aspect ratio (similar cross-sectional area) increases the range of flow rates within the monodisperse droplet producing regime (32-outlet channels and dispersed phase is 10 wt% PEG-NB, n = 1 for all data collected except h/w = 1/6 data, n = 3). A logistic regression model is applied to the collected experimental data to estimate the PDI transitional flow velocity (presented in Figure S4A).

Typically, step-emulsion systems report two flow regimes often described as stable (dripping) and non-stable (jetting) droplet generation. The separation has primarily been attributed to inertial forces (i.e., flow rate) exceeding interfacial tension^[17]^ or a critical capillary number. However, with a constant viscosity and interfacial tension, a critical flow velocity is often reported.^[19,20]^ Operating below the critical flow velocity generates monodisperse droplet populations and does not impact droplet size; however, surpassing the critical flow velocity, a jetting regime is entered where droplet dispersity and mean size increase drastically.^[19]^ Flow velocities higher than the critical flow velocity result in outer-channel isotropic droplet expansion, preventing backflow within the microchannel that produces the inter-channel pinching required for the dripping regime.^[18]^ In addition to flow impacts on stable production, aspect ratio of the outlet channel and fluid properties have also been reported to alter the ability to produce droplets in the dripping regime.^[18,19]^

A clear transition of both increasing droplet sizes and polydispersity was observed for an aspect ratio of 1/6 (h/w) when increasing flow rates (**Figure 2B**). Other buoyancy-cleared step-emulsion systems have also described a transition from uniform to non-uniform droplet production above some critical flow velocity. However, unlike previously reported results which had a drastic change in dispersity,^[15]^ we find a gradual increase in droplet variation (shown in Figure 2B right panel). In Figure 2B, droplet sizes are consistent for the first three tested flow rates of 0.5, 1, and 1.5 mL hr^−1^ (equivalent to per channel outlet flow velocities of 1.7, 3.4, and 5.0 mm sec^−1^; corresponding flow velocities are reported in **Table S1**). To further characterize this change in size and standard deviation, we use polydispersity index (PDI):

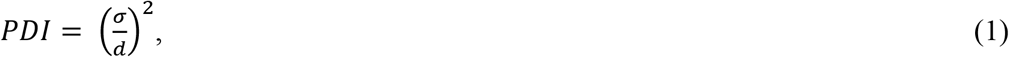

where *σ* is the standard deviation of the droplet diameter and *d* is mean droplet diameter. Monodisperse populations are defined by a PDI < 0.1 and polydisperse by a PDI > 0.1.^[31]^ Unlike traditional dripping/jetting characterizations that evaluate active droplet production in a limited field of view near the outlet channels (and thus are able to report when a critical flow velocity is reached), we sample from the entire droplet population generated after infusion of the gel precursor solution. This approach assesses an acceptable dispersity limit regardless of whether the system is operating in a dripping or jetting regime, and therefore, we report a “transitional flow velocity” (not a critical flow velocity which indicates dripping/jetting) to indicate this transition from monodisperse to polydisperse productions. Using PDI, we segment the data into mono- and polydisperse droplet production regimes, which enables us to define the transitional flow velocity, and the corresponding flow rate or throughput limitations of the system.

Across all channel geometries with the same cross-sectional area but varied aspect ratio, increasing flow rates led to both an increased mean droplet size and distribution (**Figure S3**). After assessing PDI across these devices (Figure 2C), lower aspect ratios exhibited a more clearly defined flow rate window for monodisperse production (corresponding to the tail of the sigmoid trend) while higher aspect ratios showed a more gradual increase in PDI with increasing flow velocity. We further quantified this trend using logistical regression to estimate the transitional flow velocity from mono-to-polydisperse droplet production (**Figure S4A** and **Table S2**). Lower aspect ratio devices also had lower transitional flow velocities before polydisperse droplet production. Over the flow velocities evaluated, *h/w* aspect ratios of 1/12 and 1/6 present an S-shaped dependence of PDI on flow velocity. In contrast, aspect ratios of 1/3 and greater appear to follow a similar sigmoidal relationship but with slopes and midpoints sufficiently shifted so that the S-curve is not entirely captured within the tested flow rate ranges. PDI trends for increasing flow velocity resulted in different points of transition before producing polydisperse droplet populations (**Figure 2C**). Specifically, devices with larger *h/w* aspect ratios sustain higher flow velocities while maintaining monodisperse droplet generation.

#### 2.1.2 Aspect ratio influences droplet size in channels with similar cross-sectional area outlet geometries

We investigated the effect of channel aspect ratio on the resulting droplet sizes produced by evaluating the lowest tested flow rate, 0.5 mL hr^−1^ (corresponding to the lowest flow velocities in Figure 2C). This flow rate was selected as it corresponds to the most “stable” (lowest PDI) droplet producing flow velocities across all devices, eliminating any flow-induced dispersity effects across channels that may be close to the transitional flow velocity. Furthermore, we verified that the droplet production at these flow rates mostly resulted in the dripping regime for all aspect ratios (except 1/1 and 2/1) using high-speed imaging (Figure S4B). Mean monodisperse droplet sizes (at 0.5 mL hr^−1^) increase with aspect ratio (**Figure 3**) until h/w aspect ratio reaches 1 before decreasing. These results indicate that the droplet size is not height dependent, but rather dependent on the smallest channel dimension. Normalized droplet diameter (*d*_*droplet*_*/d*_*avg droplet*_) distributions at similar flow rates are consistent for all aspect ratios (Figure 3C), indicating that the spread of the droplets is similar.

**Figure 3.**
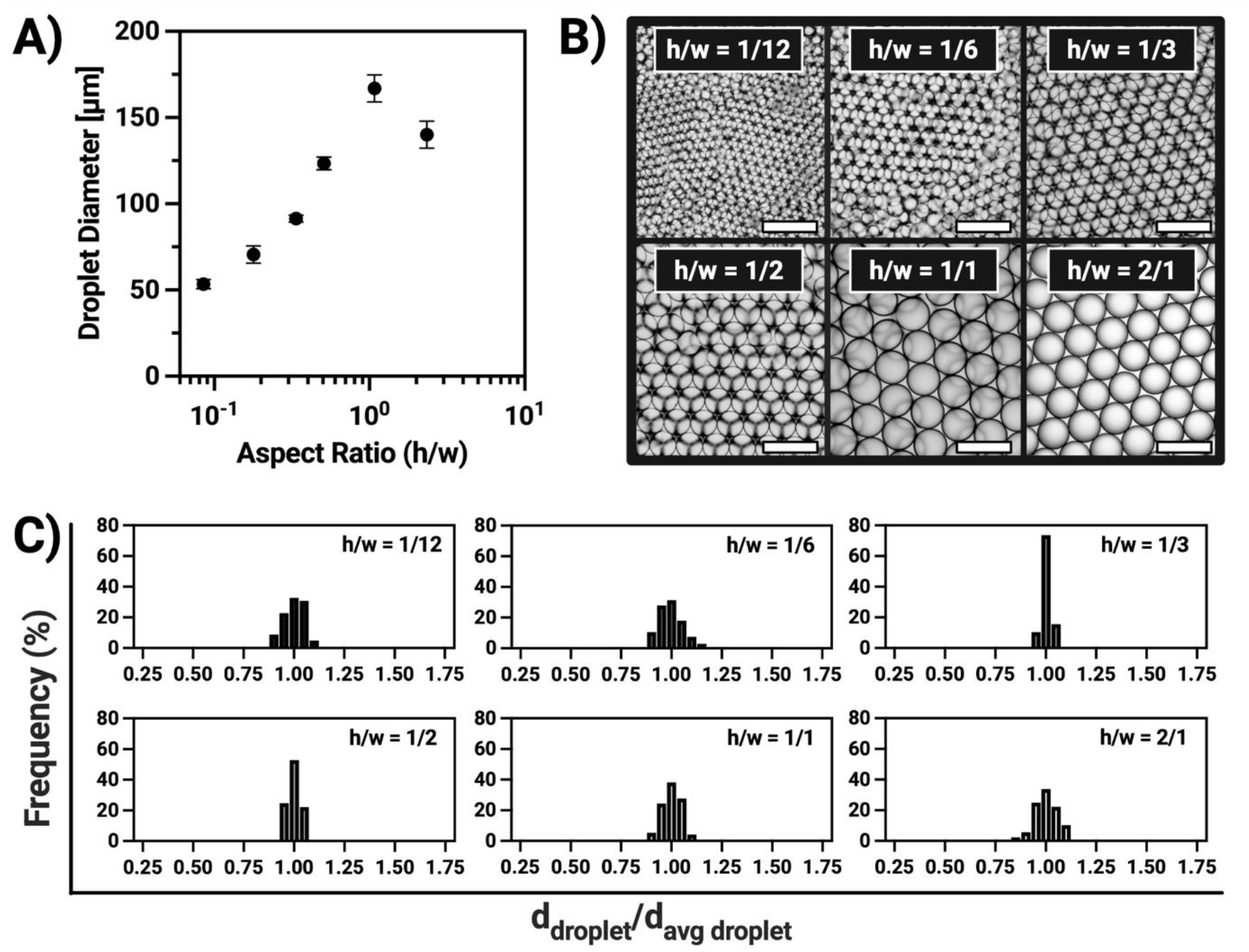
At a similar cross-sectional area, droplet size in the dripping regime is dependent on the aspect ratio of the channel. By operating the device at 0.5 mL hr^−1^ (most monodisperse flow rate tested), we identified that droplet diameter is largest when aspect ratio is close to 1. Evaluation of the different aspect ratio droplet generation used 32-outlet channel devices and a dispersed phase of 10 wt% PEG-NB. A) Average droplet diameter plotted as a function of channel aspect ratio (height/width); error bars are standard deviation of the distribution of droplet sizes. B) Brightfield images from monodisperse droplet generation at 0.5 mL hr^−1^ for varying aspect ratios and similar cross-sectional area; scale bars are all 250 µm. C) Frequency distributions of the normalized droplet diameter for each device type show narrow distributions across all aspect ratios tested; device was operated at 0.5 mL hr^−1^.

Previous work has demonstrated that droplets formed in the dripping regime had diameters (*d*) which scaled with outlet channel height (vertical-oriented outlet channels), reporting *d* ∼ 4h^[18]^ (and similarly *d* ∼ 4.12h in related experiments^[15]^). Our measured droplet sizes from our aspect ratio assessment (Figure 3A) follow a comparable scaling, with *d* ∼ 3 to 4h across almost all devices (Figure S4B). Importantly, our outlet geometry differs from the cited work in that the droplet expansion into the reservoir in our device occurs both planarly and vertically up while the bottom boundary remains fixed, so height and width are not interchangeable. Consistent with this directionality, the 2/1 (h/w) device deviates from height-based scaling, and to maintain a dimensionless droplet diameter relationship of 3 to 4, width-based scaling was used (Figure S4B). Together, these results suggest that in our architecture, the gel precursor droplet size is governed primarily by the most constraining (smallest) outlet dimension (whether that be height or width), rather than by aspect ratio alone.

Additionally, previous work identified a critical channel aspect ratio of ∼ 2.6 (vertical-oriented outlet channels) as the transition between dripping (which happens when 1/2.6 > h/w and when h/w > 2.6/1) and jetting (when 1/2.6 < h/w < 2.6/1).^[18]^ Interestingly, we observed dripping at an aspect ratio of 1/2 (at 0.5 mL hr^−1^) which was supported by high-speed image capture with visual confirmation of necking (Figure S4C). Although we did not map dripping/jetting boundaries across flow rate, high-speed imaging at 0.5 mL hr^−1^ confirmed dripping for all tested geometries except the 1/1 (isotropic expansion, a hallmark of jetting) and 2/1 (h/w) device (for which dripping/jetting could not be conclusively assessed due to limitations in the camera orientation, Figure S4D).

### 2.3 Producing distinct droplet diameter sizes based on outlet channel dimensions

One of the goals of our work was to establish a guide for other researchers to easily adopt this technology with predictable droplet size outputs based on device design parameters. Accordingly, we evaluated the impact of outlet channel geometries on gel precursor droplet sizes to define a relationship between droplet size and channel dimensions. We assessed droplet generators with channels of varying heights, widths, and aspect ratios and evaluated their corresponding droplet populations which ranged from a mean droplet diameter of 54 µm to 559 µm (**Figure 4A**). Droplet generation from each device was completed in the monodisperse regime and presented from smallest to largest mean droplet diameter. Droplet generators assessed either had 32- or 512-outlet channels. The increase of parallelized outlet channels enabled us to increase our input flow rates and consequently achieve faster droplet generation. To evaluate geometric parameters that correlated to droplet sizes, we plotted various geometric parameters and combinations of parameters against mean droplet diameter (**Table S3** and **Figure S5**). We found significant correlations (R^2^ > 0.7) within the data sets we tested (Figure S5). In step-emulsion droplet generators where the dispersed phase flows and exits the channels vertically, height is considered the smallest dimension and the primary factor determining droplet size.^[15,19]^ Due to the orientation of our lateral ledge emulsion droplet generator, the smallest dimension could be height or width, and among the geometric parameters we tested for correlation to predict droplet size (Figure S5), we confirmed “smallest dimension” as the defining parameter of droplet size (Figure 4B) with an R^2^ = 0.97. It is important to note that although there is a high correlation for “smallest dimension”, fewer geometries with an aspect ratio above h/w = 1 were tested due to the increased difficulty of fabricating and maintaining high aspect ratio outlet channels. With this in mind, the smallest dimension is still most correlated with a broader range of aspect ratios. When considering the geometries we tested, when aspect ratio is less than 1 (i.e., height is the smallest dimension), we also observe a similar trend (R^2^ = 0.97). While the smallest dimension of the outlet channel should provide a good indication of the expected droplet size, to increase the certainty that this is a salient predictor of droplet diameter with our step-emulsion device, a larger sample set of > 1 aspect ratios would need to be evaluated.

**Figure 4.**
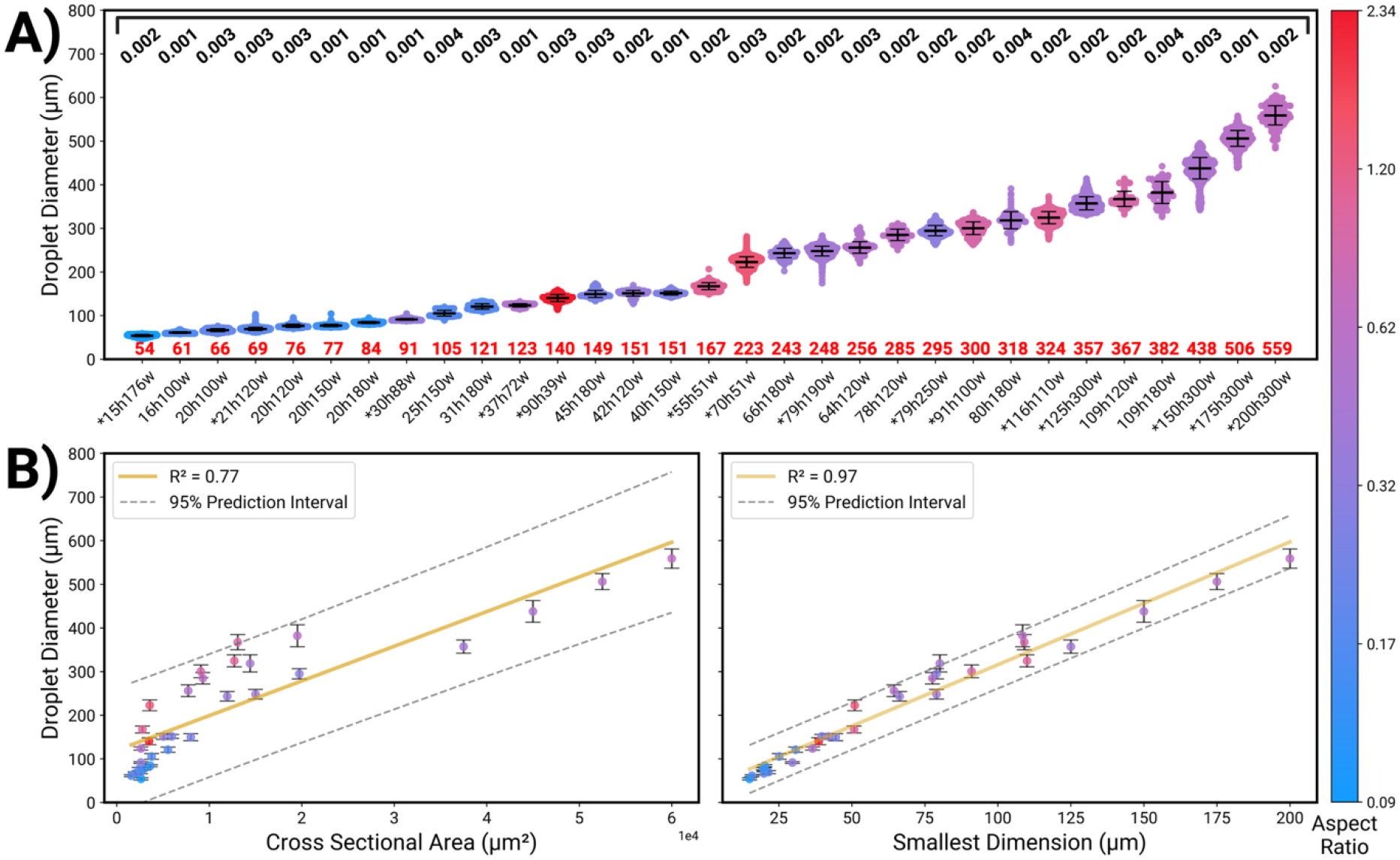
We evaluated a broad set of droplet generators and identified smallest dimension (either height or width) as the parameter most correlated to gel precursor droplet size in our system. A) Monodisperse droplet populations (dispersed phase is 10 wt% PEG-NB) produced from each tested geometry are shown, with polydispersity index (PDI) reported above the scatter dot plots (in black text) and average droplet diameter (µm) below (in red text). Channel dimensions are given in the x-axis labels, with channel height and then width reported (i.e., 15h176w corresponds to channel dimensions of 15 µm height and 176 µm width). Dataset includes droplet populations from both 32- and 512-outlet channel devices. All devices tested as 32-outlet channels are denoted with an asterisk in front of the height and width dimensions label. B) The correlation between droplet diameter and cross-sectional area (left) or smallest dimension (right) is visualized here with a 95% prediction interval (see Figure S5 for additional geometric parameters evaluated). While cross-section correlates with droplet diameter, smallest dimension exhibits the strongest association, underscoring its importance in designing systems for targeted droplet sizes. Aspect ratios for all data are reported throughout the figure as a heat map (on right). Error bars are standard deviations.

### 2.4 Robust gel precursor droplet generation

#### 2.4.1 Material and user independence

Most studies that present droplet generation systems for microgel production traditionally validate the device with a single gel precursor droplet solution. Our step-emulsion droplet generator should produce droplets through Laplace pressure induced snap-off^[32]^ and function across a range of viscosities and with relatively similar flow rates. We wanted to verify that our step-emulsion system indeed could be applied to gel precursor solutions of different backbone compositions and gelation mechanisms (**Table S4**). Accordingly, we evaluated material compatibility with our system by testing various gel precursors (PEG, HA, alginate, gelatin, trehalose) with various gelation mechanisms (ionic, Michael addition, thiol-ene click reaction, strain-promoted azide–alkyne cycloaddition, free-radical photopolymerization) using our 512-outlet channel device with channel dimensions of 20 µm height x 120 µm width (20h120w). In these tests, the gel precursor solution was infused through our system at 10 mL hr^−1^ (2.25 mm sec^−1^ per channel outlet velocity, **Figure 5A**, top panel). Across all materials at constant flow (10 mL hr^−1^), droplet populations were monodisperse (PDI < 0.1) and all droplet diameter means are within 10% of each other. We attribute the slight changes in mean droplet diameter to deviations in material properties (density, viscosity, and surface/interfacial tension) or minor variability in device fabrication.

**Figure 5.**
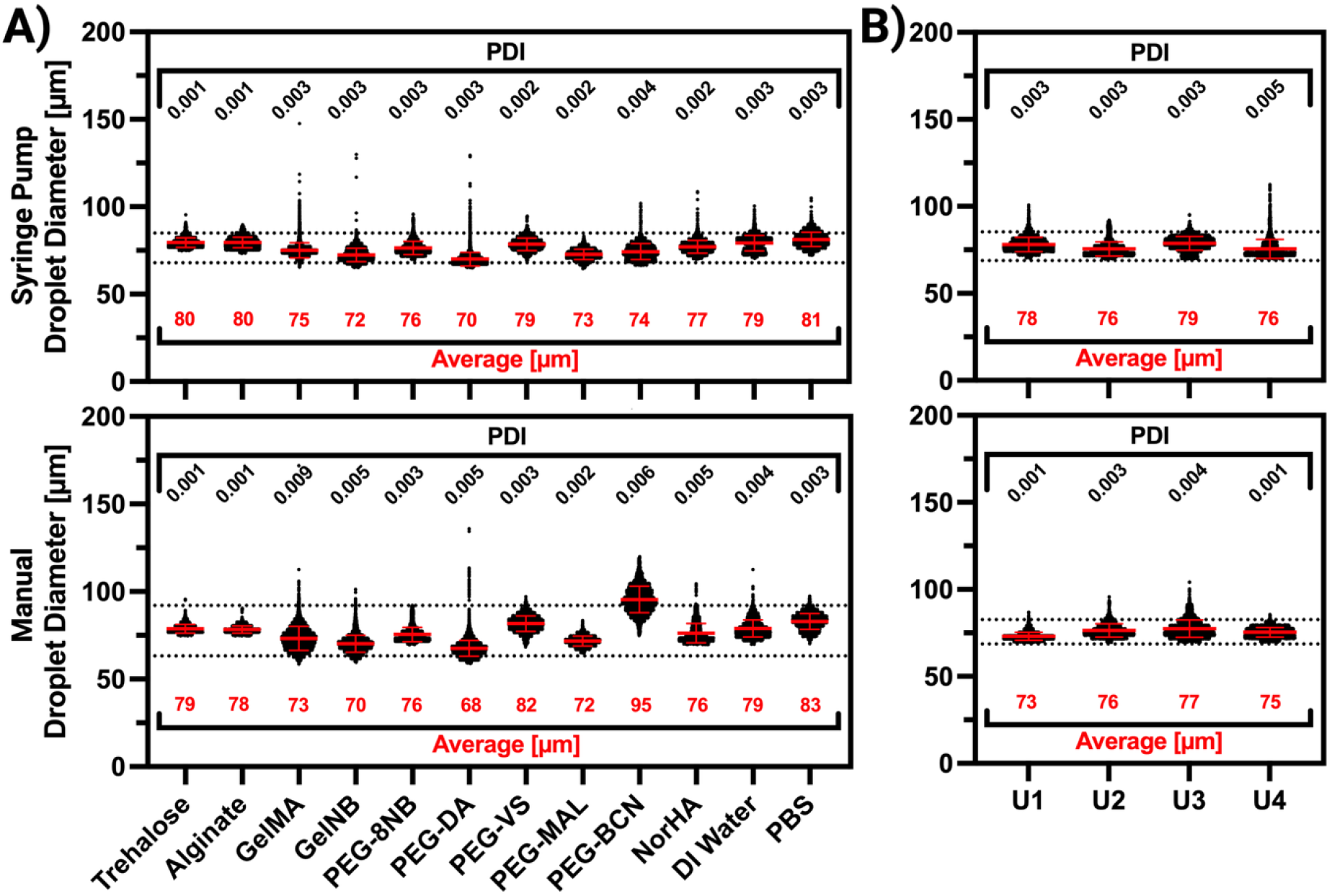
In the stable droplet producing regime, droplet generation is relatively independent of flow rate, user, and gel formulation. Due to the flow velocity independence, the system can be operated by either a syringe pump (top panel) or manual injection (bottom panel). A) Droplet populations using various biomaterial gel precursor solutions are presented, where the dashed lines in the plots indicate the tolerance interval encompassing 90% of the population with 95% confidence (67.9 to 84.9 µm for 10 mL hr^−1^ and 63.2 to 92.0 µm for manual injection). B) The device demonstrates user independence, consistently producing similar droplet populations across different users through syringe pump (top panel) or manual (bottom panel) injection of the gel precursor. Dashed lines indicate the tolerance interval encompassing 90% of the population with 95% confidence (68.6 to 82.6 µm for 10 mL hr^−1^, 68.8 to 85.3 µm for manual). Schematics above compare 10 mL hr^−1^ operation and schematics below compare manual-driven operation. For all plots, PDI is reported above the scatter dot plots (in black text) and the average droplet diameter (µm) below the scatter dot plots (in red text). Error bars within the scatter dot plots represent standard deviation.

Due to the relative insensitivity of PDI to flow velocity changes while maintaining device operation under the critical flow velocity (Figure 2B), we evaluated manual injection of gel precursors into our device as a simple droplet generation technique. As long as the user is pushing at a flow velocity below the transitional flow velocity, the device should still yield a monodispersed droplet population, despite the non-continuous input flow rate or flow velocity in the channels. The manual droplet populations favorably resembled their constant (10 mL hr^−1^) counter parts but with slightly larger deviations in mean droplet size. One exception was seen with PEG-tetrabicyclononyne (PEG-BCN), a SPAAC-based chemistry that occurs rapidly at elevated temperature. We attribute this to the inconsistent nature of the supplied flow and the conditions in which it was run as heat was inadvertently supplied from the hand to the syringe containing the prepolymer during injection (PEG-BCN was infused into the device in a cold room due to the rapid nature of the polymerization reaction). The resulting droplet populations of the various biomaterials sat within a 90% tolerance interval (capturing 90% of the population with 95% confidence, determined from power-based binned averaging of droplet diameter distributions) from 67.9 to 84.9 µm for 10 mL hr^−1^, and 63.2 to 92.0 µm for manual injection demonstrating compatibility across multiple materials. It is useful to note that by excluding the manual injection of PEG-BCN from the 90% tolerance interval (with 95% confidence), the manual injection tolerance interval contracts (65.1 to 86.8 µm) – falling strikingly close to the 10 mL hr^−1^ syringe pump injection tolerance interval.

One of our major goals was to develop a simple, single-step platform to produce monodisperse microgels across different biomaterials, but also to enable easy adoption by researchers outside the microfluidic space. To assess the robustness and versatility across users of different experience levels with both this device and more generally with microfluidics, we employed a collaborative testing approach with researchers across different research labs (Figure 5B). Here, users were given the same gel precursor and operated the device both manually and with a syringe pump. By infusing slowly, users consistently produced monodisperse droplets through manual injections regardless of their experience with microfluidics.

#### 2.4.2 Capillary number as an estimator for assessing increasingly viscous solutions

Monodisperse droplet populations were achieved across all tested biomaterial precursor solutions up to 10 mL hr^−1^. For a fixed outlet channel geometry, the transition from dripping to jetting is primarily governed by a critical capillary number.^[19,20]^ Therefore, we used a critical capillary number to estimate at what flow velocity (and corresponding flow rate) the droplet production would transition from a stable dripping to a jetting regime. Using a critical capillary number (Ca = 0.014) previously reported by Montessori et al. for all materials^[20]^, we estimated the transitional flow velocities between the two regimes and operated our device below these maximum dripping flow velocities. We assessed our device’s compatibility across solutions of varying viscosity by evaluating solutions with viscosities ranging from 1 to 669 mPa s (viscosities of gel precursors tested in Figure 5 varied between 1 and 40 mPa s). To achieve large changes of viscosity with a common biomaterial, we used alginate concentrations from 2 to 5 wt%. A list of the viscosities and interfacial tension between the aqueous droplet solutions - water, PEG-NB, and alginate at 2, 3, 4, and 5 wt% - and the fluorinated oil are provided in **Table S5**. To test varying viscosities, we used our 20 µm height x 120 µm wide emulsion device for droplet generation. By flowing under the estimated maximum dripping flow velocity, we produced monodisperse droplet populations independent of their respective viscosities (**Figure 6**).

**Figure 6.**
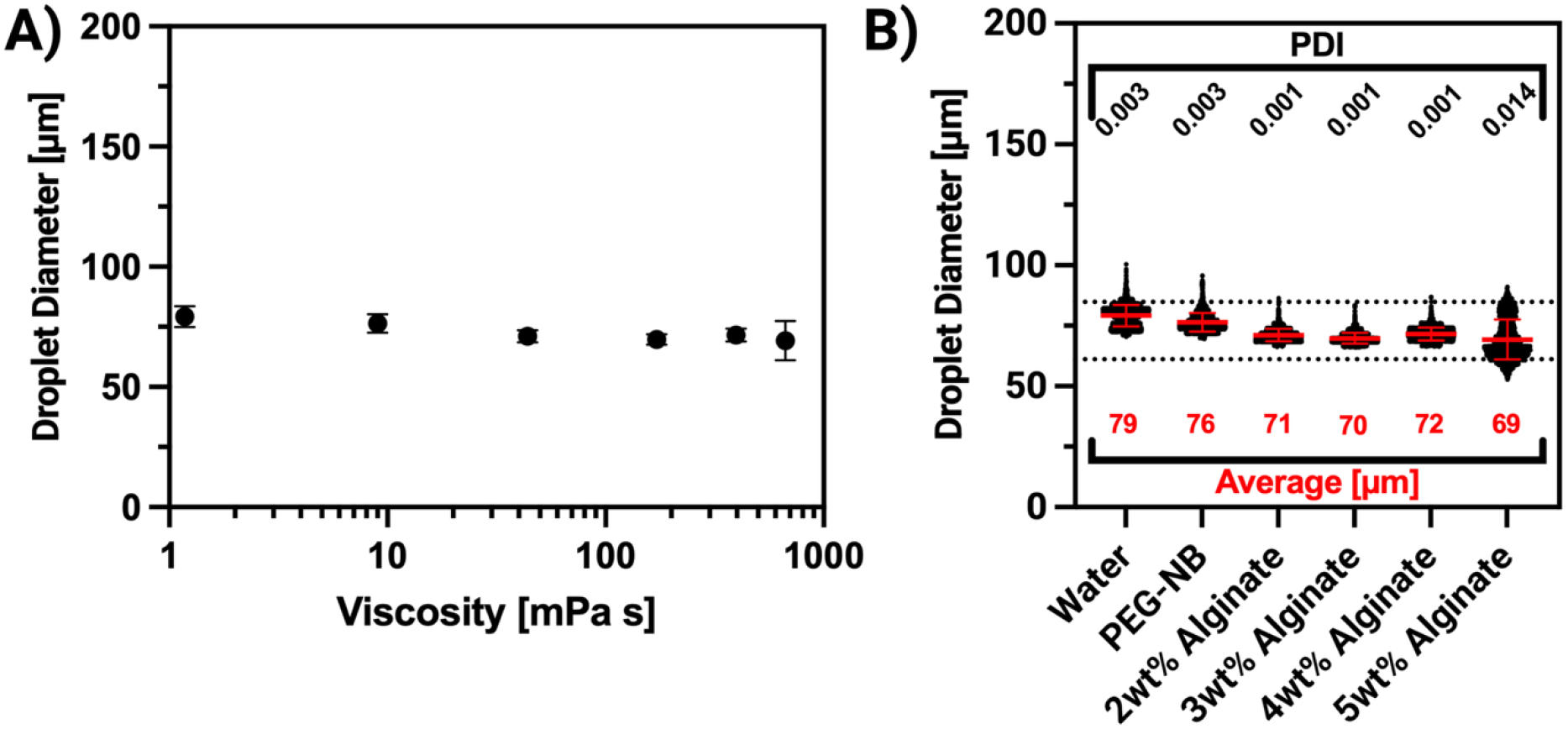
The step-emulsion device produced monodisperse droplet populations, with similar mean droplet diameter across solutions with viscosity ranging from 1 to 669 mPa s. A) When operating below the estimated capillary number, droplet diameter remained independent of viscosity. B) Corresponding scatter dot plots of data presented in 6A reporting droplet size distributions and ordered by increasing fluid viscosity from left to right (dashed lines indicate the tolerance interval for 90% of the population with 95% confidence, 61.1 to 84.8 µm). Polydispersity index (PDI) is reported above the scatter dot plots (in black text) and average droplet size (µm) below (in red text). Error bars in both plots represent standard deviation.

While we had to use reduced flow velocities for the different alginate solutions (2, 3, 4, 5 wt%), we were able to operate the system at 4, 2, 0.5, and 0.3 mL hr^−1^ (Table S5), respectively. To the best of our knowledge, uniform droplet generation of 5 wt% alginate (with viscosity of 669 mPa s) has not been presented in microfluidic droplet generators at these sizes and production rates. Additionally, all the alginate droplet populations fall within a 90% tolerance interval (95% confidence) of 61.1 to 84.8 µm, which is similar to the 90% tolerance interval reported in **Figure 5A**, when using a syringe pump (68.6 to 82.6 µm). The shift with the alginate is largely due to the 5 wt% alginate gel having a larger droplet spread, which may be due to the difficulty of mixing and loading alginate into the syringe with no microscopic air bubbles. This variation may also be from the inherent variability of the syringe pump operation and the associated instantaneous flow rate fluctuations of most syringe pumps^[33]^ briefly surpassing the maximum dripping flow velocity. Critically, these tests demonstrate that we can achieve relatively high throughput monodisperse droplet generation even when using materials with differing viscosities and interfacial tensions, highlighting the robustness of our droplet production platform.

## 3. Conclusion

In this work, we generated a platform for simplified, high throughput, and uniform droplet generation to streamline microgel production workflows. Conventional methods for producing uniform microgels often require relatively complicated experimental setups, particularly for non-experienced microfluidicists, and require extensive calibration to generate uniform gel precursor droplets of discrete size, while also being constrained by limited production rates. To address this, we developed a device that leverages parallelized step-emulsion microchannels which, when operated in a stable producing regime below a critical flow velocity, reliably produce droplet populations for monodispersed microgel and granular hydrogel generation. We characterized this flow independence in the stable regime across all devices and used PDI as a metric to assess droplet uniformity as a function of flow velocity.

We probed the extent of the monodisperse flow regime in devices with similar cross-sectional area but varying aspect ratios, finding that as aspect ratio increases, the flow range for achieving monodispersity increases as well. Our experimental data on aspect ratio effects aligns with prior findings. Analysis of all collected data confirms smallest dimension as the primary geometric parameter defining droplet size in our system. We further demonstrated the robustness of our platform by generating monodisperse droplets across various commonly used biomaterials with diverse fluid properties and validated its ease of use through multi-user testing. We have developed a droplet generation system that can be operated in a plug-and-play manner, with multiple biomaterial gel precursors, producing highly consistent and monodispersed droplets enabling scalable microgel production for a broad range of applications.

## 4. Experimental Section/Methods

### Device Fabrication

Molds for devices with heights 125 µm or higher were designed in Fusion 360™ (available in supplemental materials) and fabricated using a 3D printer (J750 Verowhite, Stratasys, Eden Prairie, MN). All other molds for devices were designed in Autocad™ and fabricated using standard SU-8 photolithography (SU-8 2000, Kayaku Advanced Materials, Westborough, MA) protocols from the manufacturer and surface treated with 1H,1H,2H,2H-perfluorooctyltrichlorosilane, 97% (#AAL1660609, Fisher Scientific). Files for examples of 3D printed molds (.stl file) and SU8 molds (.dwg) are provided in supplemental files for easier access to device fabrication. Molds generated using SU8 photolithography were 2-layer molds, using 2 photomasks. The 1^st^ device layer contained the droplet generation channel reported on heights and widths. The 2^nd^ layer was to create a more prominent edge at the channel outlets, and to serve as a guide for optimal cutting of the device. The 2^nd^ layer also increased the height of channels near the device inlet to reduce the total device resistance and associated pressure needed to provide the desired flowrates. When generating the 2^nd^ layer, the 2^nd^ layer height was at least 4 times the height of the 1^st^ layer. PDMS devices were cast on the molds using Sylgard 184 poly(dimethylsiloxane) (PDMS; Dow, Midland, MI), which was prepared at a 10:1 mixture and degassed. PDMS was poured into molds and cured at 60 °C for at least 4 hrs in a convection oven. Uncured PDMS was also used to spin coat glass slides at a thickness of 40 µm and cured at 120 °C in a convection oven for at least 15 min. After curing, PDMS device layers were removed from the mold, and inlets were made using a 2 mm biopsy punch (#33-31-P, Integra Lifesciences, Princeton, NJ). Outlet channel heights were measured from the delaminated PDMS using an Olympus OLS 4000 LEXT. The delaminated PDMS was carefully and precisely cut into individual devices using a flat razor blade (11-515, Stanley, Seattle, WA). Note: Precise cutting at the outlet channel edge is crucial as non-ideal cuts in this step can heavily impact the device’s ability to produce monodisperse droplets. After verifying the quality of the cut, each device was then plasma bonded onto PDMS coated glass slides. These assemblies were placed in a 60 °C oven for at least 24 hrs to regain the channel’s hydrophobicity. The glass slide was then cut using a glass cutter so that the device may fit into a 60 mm petri dish. Devices underwent final inspection using a microscope to assess devices for non-optimal bonding and edge of device delamination and to further review cuts made at the outlet channel. The device was adhered to the petri dish using double sided tape (Scotch Double Sided Tape, 3M, Maplewood, MN) and stored under vacuum in a vacuum desiccator until use.

### Droplet Generation

Gel precursor was prepared and loaded into a 1 mL Luer-lock syringe (#309628, Becton Dickinson, Franklin Lakes, NJ). The syringe was turned with the outlet facing up and any visible air bubbles were removed by tapping the syringe. A 14-gauge Luer-lock blunt tip (#JG14, Jensen Global, Santa Barbara, CA) was attached to the syringe and inserted into 1/16” ID tubing (#MFLX06422-02, Avantor, Radnor, PA). A cannula taken from a 14 gauge blunt tip was placed into the opposite end of the tubing and the entire assembly was placed on a syringe pump. The petri dish containing the device was taken out of the vacuum desiccator and filled with 0.4% (w/w) Pico-Surf® (#C022, SphereBio, Cambridge, UK) in Novec™ 7500 (3M, Maplewood, MN) until the oil submerged the device. Air bubbles in the device were removed by pipetting oil into the device inlet to flush the channels and any residual air bubbles appeared to be absorbed by the oxygen-deprived PDMS. The cannula was placed into the device and gel precursor was infused with either a syringe pump (Fusion series, Chemyx, Stafford, TX) or manually (both with 300µL of solution). Post droplet generation, images were taken in the 60 mm petri dish to decrease overlap of droplets for improved image analysis. Images were analyzed using MATLAB script as previously described.^[34]^ After droplet generation and image collection (PEG-BCN gelation was immediate and therefore, the images are of microgels), the droplets were gelled using the methods described for each material and poured into a 50 mL conical tube for collection. A detailed protocol for droplet generation is included as supplemental documentation and for microgel purification (as an additional document) we have applied to the following microgel types as a resource for the broader scientific community. These protocols were provided to users in different research labs as part of the collaborative assessment. The instructions given to users to generate Figure 5B are also included as part of supplemental documents.

### PEG-NB Microgel Fabrication

8-arm poly(ethylene glycol)-norbornene (PEG-NB, M_n_ ∼ 40,000 Da, Creative PEGWorks, Chapel Hill, NC) and poly(ethylene glycol) dithiol (PEGDT; M_n_ ∼ 3,505 Da, JenKem Technology USA, Plano, TX) were purchased already synthesized from commercial sources. Lithium phenyl-2,4,6-trimethylbenzoylphosphinate (LAP; Sigma-Aldrich) and tris(2-carboxyethyl)phosphine hydrochloride (TCEP; Sigma-Aldrich) were used as the photoinitiator and reducing agent, respectively. PEG-NB precursor was prepared in phosphate-buffered saline (PBS; pH 7.4) to a final composition of 10 wt% PEG-NB (2.27 mM) and a thiol:ene ratio of 0.4:1 (3.64 mM PEGDT, 2.20 mM LAP, and 0.73 mM TCEP). The precursor solution was then loaded into a syringe and infused into the device as reported. Post droplet generation, the droplets underwent gelation by exposure to 365 nm UV light (200 mW/cm^2^ for 60 sec).

### Trehalose Droplet Fabrication

Trehalose (Pfanstiehl, Waukegan, IL) was prepared at 1 M in deionized water and infused into the 20h120w device at 10 mL hr^−1^ and manually.

### Alginate Microgel Fabrication

Alginate precursor was prepared as previously described by Utech et al.^[35]^ Briefly, a calcium-EDTA solution was prepared by mixing 100 mM calcium chloride (#BP510, Fisher Scientific) and 100 mM disodium-EDTA (#E5134, Sigma-Aldrich) at a ratio of 1:1 and pH adjusted to 7.2 using 1 M sodium hydroxide (#221465, Sigma-Aldrich). Sodium alginate (#W201502, Sigma-Aldrich) was then dissolved in the calcium-EDTA solution at 2, 3, 4, and 5 wt% and loaded into a 1 mL syringe. The alginate solution was infused into the 20h120w device at 10 mL hr^−1^ and manually. Post droplet generation, the droplets were introduced to a 0.05 vol% acetic acid (#A38212, Fisher Scientific) in Novec™ 7500 solution to initiate gelation.

### GelMA Microgel Fabrication

Gelatin-methacrylate (GelMA; 300 bloom, DS 53% Allevi Inc by 3D Systems, Philadelphia, PA) was lyophilized and reconstituted in phosphate buffered saline (PBS; pH = 7.4) at 20% (w/v). The prepolymer solution was placed on a heat block to warm the solution to 37 ℃, and GelMA hydrogel precursor solutions were made with 5% (w/v) GelMA prepolymer solution and 1 mM LAP in PBS at 37 ℃. GelMA precursor was infused into the 20h120w device at 10 mL hr^−1^ and manually. After droplet generation was complete, droplets were exposed to 365 nm UV light (5 mW/cm^2^ for 180 sec).

### Synthesized Compounds

Poly(ethylene glycol) tetrabicyclononyne (PEG-tetraBCN, M_n_ ∼ 20,000 Da) and gelatin-norbornene (Gel-NB) were synthesized as previously reported.^[36,37]^ Gel-NB degree of substitution (DS) was determined by ^1^H-NMR (Bruker Avance-I Series 300.10 MHz), using the method described in the literature.^[37]^ Norbornene-modified Hyaluronic Acid (NorHA) was synthesized following the method reported by Plaster et al.^[38]^ Briefly, Sodium Hyaluronate (M_n_ ∼ 90,000 Da, lot# 030207, Lifecore Biomedical) was solubilized in 2-(N-morpholino) ethanesulfonic acid buffer (MES; pH 5.5; lot#SLC06070, Sigma-Aldrich) and later reacted with 4-(4,6-dimethoxy-1,3,5-triazin-2-yl)-4-methylmorpholinium chloride (DMTMM; lot# WUFCH-TY TCI America, Portland, OR) and 5-norbornene-2-methylamine (lot# F5JHF-NL, TCI America, Portland, OR). Saturated sodium chloride was added to the mixture prior to precipitation with 200 proof ethanol dropwise. The final precipitate was collected using vacuum filtration and resolubilized in deionized water. The final product was then dialyzed in DI water for 3 days using 6-8,000 Da dialysis tubing (lot#20100189, Repligen, Boston, MA) with the final product frozen and lyophilized. The lyophilized product was stored at -20 °C. NorHA degree of substitution was measured using ^1^H-NMR (Varian Series 400MHz), with NorHA dissolved at 10 mg/mL in deuterium oxide.

### Gel-NB Microgel Fabrication

Gel-NB was dissolved in PBS at 10% (w/v) to form a prepolymer solution. The prepolymer solution was set on a heat block to warm to 37 ℃, and Gel-NB hydrogel precursor solutions were made with 5% (w/v) Gel-NB prepolymer solution and a thiol:ene ratio of 1:1, (5.775 mM dithiothreitol (DTT; Dithiothreitol, 98.0+%, #D107125G, TCI America, Portland, OR) and 1 mM LAP) in PBS at 37 ℃. Gel-NB precursor was infused into the 20h120w device at 10 mL hr^−1^ and manually. After droplet generation was complete, droplets were exposed to 365 nm UV light (5 mW/cm^2^ for 180 sec).

### PEGDA-8000 Microgel Fabrication

Poly(ethylene glycol) diacrylate (PEGDA-8000; M_n_ ∼ 8,000 Da, Catalog #046801.03, Thermo Scientific Chemicals, Haverhill, MA) and 2,4,6-trimethylbenzoylphosphinate (LAP; Catalog #L0290, TCI America, Portland, OR) were purchased from commercial sources. To form PEGDA-8000 microgels, hydrogel precursor solutions were made with 10% (w/v) PEGDA-8000 and 1 mM LAP suspended in deionized water at room temperature. PEGDA-8000 precursor was infused into the 20h120w device at 10 mL hr^−1^ and manually. After droplet generation was complete, droplets were exposed to 365 nm UV light (5 mW/cm^2^ for 180 sec).

### PEG-tetraBCN Microgel Fabrication

To decrease the gelation rate between -BCN in PEG-tetraBCN and -azide in crosslinker, all material preparations and microgel fabrications were performed in a cold room at 4 ℃. Synthesized PEG-tetraBCN (M_n_ ∼ 20,000 Da) was resuspended in PBS to form a 10 mM stock solution. The PEG-tetraBCN stock solution was further diluted with PBS, and (triethylene glycol) diazide (TEG-diazide; Catalog #207, Lumiprobe Corporation, Westminster, MD) was added to the mixture at 4 ℃ so that the final hydrogel precursor solution was 4 mM PEG-tetraBCN: 8 mM TEG-diazide. To prevent early gelation, the mixed solution was immediately drawn into a 1-mL Luer-lock syringe for manual and 10 mL hr^−1^ syringe-pump injections with 20h120w devices, as previously stated.

### NorHA Microgel Fabrication

Norbornene-modified hyaluronic acid (NorHA) (%DS 31.5) was first dissolved in phosphate buffered saline (PBS; pH = 7.4) with continuous mixing and heating at 37 ℃ on a vortex mixer at a solution concentration of 0.5 mM. The gel precursor solution was then made at concentrations of 2% (w/v) NorHA (0.21 mM), 2.33 mM PEGDT, 1.12 mM LAP, 0.23 mM TCEP, and 0.01 mM Dimethylformamide (DMF) dissolved in PBS at room temperature. NorHA precursor was infused into the 20h120w device at 10 mL hr^−1^ and manually. After droplet generation was complete, droplets were exposed to 365 nm UV light. *PEG-MAL Microgel Fabrication:* 8-arm poly(ethylene glycol) maleimide (PEG-MAL, M_n_ ∼ 40,000 Da, >90% purity, JenKem Technology USA, Plano, TX) was pre-functionalized with RGD peptide (Ac-GCGYGRGDSGP, MW = 1067.10 g/mol, >85% purity, GenScript USA Piscataway, NJ) by suspending PEG-MAL and RGD in HEPES (Isotonic HEPES buffer, 50 mM, pH 7.4), mixing thoroughly, and allowing the reaction to proceed for 15 min at 20 °C. The pH of the PEG-MAL/RGD mixture was then lowered with HCl. Separately, YKNS peptide (Ac-GCYK↓NSGCYK↓NSCG, MW = 1525.69 g/mol, >90% purity, GenScript USA Piscataway, NJ) was prepared by suspending in TCEP and subsequently added to the PEG-MAL/RGD/HCl mixture. The final microgel precursor solution consisted of 10 wt% PEG-MAL, 1 mM RGD, and 2.77 mM YKNS (corresponding to a 16:7 stoichiometric ratio of maleimide to thiol between PEG-MAL and YKNS). The precursor was infused into the 20h120w device at 10 mL hr^−1^ and manually. After droplet generation was complete, the droplets were allowed to react overnight at room temperature.

### PEG-VS Microgel Fabrication

8-arm poly(ethylene glycol) vinyl sulfone (PEG-VS, M_n_ ∼ 40,000 Da, >90% purity, JenKem Technology USA, Plano, TX) was suspended in isotonic HEPES buffer (pH 6.5) while YKNS was suspended in TCEP. PEG-VS and YKNS solutions were combined and mixed thoroughly. The final microgel precursor solution was composed of 5 wt% PEG-VS and 2.67 mM YKNS (corresponding to a 1:0.8 stoichiometric ratio of vinyl sulfone to thiol between PEG-VS and YKNS). The precursor was infused into the 20h120w device at 10 mL hr^−1^ and manually. After droplet generation was complete, the droplets were allowed to react overnight at room temperature.

### Viscosity Measurements

Gel precursor solutions were prepared as described above without crosslinker or photoinitiator, and viscosity measurements were performed using a Peltier double-wall concentric cylinder (Aluminum) on the Discovery HR-30 rheometer (TA Instruments, New Castle, DE). Briefly, the temperature was initially set to 25 °C, and 13 mL of polymer solution was poured into the lower cup geometry. After that, a double-gap rotor was installed with the instrument, and viscosity data was collected at a 2 mm gap over the 0.01-1000 Hz shear rate range.

### Interfacial Tension Measurements

Gel precursor solutions were prepared as described above except with PEG-NB being made without LAP. Interfacial tension measurements were performed using the pendant-drop method on a Theta Lite optical tensiometer (Biolin Scientific) and analyzed using the instrument’s built-in image-analysis software. A quartz cuvette was filled with 0.4 wt% PicoSurf in Novec 7500 sufficient to immerse the J-hook needle dispensing tip. The gel precursor solution (∼1 mL) was loaded into a plastic syringe, then fitted with a J-hook needle and dispensed into the oil phase to form pendant droplets. For each gel precursor composition, three droplets were generated and analyzed; only droplets that remained intact for at least 1 min were included. Interfacial tension values were recorded at steady state.

### High-speed Imaging

Visualization of droplet formation was performed on a Thorlabs Veneto inverted microscope equipped with a motorized stage (MLS203, Thorlabs, Newton, NJ). Samples were placed on the stage and imaged using a cavitation light path with transmitted light (trans-illumination module). Images were acquired through a 10X objective (Nikon Plan Apo λ, 10X/0.45) and recorded using a Shimadzu high-speed camera (HPV-X2, Shimadzu, Columbia, MD) controlled with HPV-X software (Shimadzu, Columbia, MD). PEG-NB precursor (prepared as described above but without LAP) was infused at 0.5 mL hr^−1^ and high-speed imaging was taken at 10 kfps for all aspect ratios except 21h120w (20kfps).

## Supporting information

CAD Files, PDF of Device Design, Text Description

Droplet Generator Operating Protocol Provided to New Users

Microgel Purification Protocol Provided to New Users

Video of Droplet Generator Producing Water droplets with Food Dye Top View

Video of Droplet Generator Producing Water droplets with Food Dye Side View

Video of Droplet Generator Producing Water droplets with Food Dye Side View 2

Video of water droplets emerging from channels

## Acknowledgements

This research was supported by graduate research fellowships to D.P.L from the National Academies of Sciences, Engineering, and Medicine through the Ford Foundation Predoctoral Fellowship and from the Rackham Graduate School at the University of Michigan through the Rackham Merit Fellowship, with additional funding provided by the Navajo Nation. This research was also supported by the NIH’s Maximizing Investigators’ Research Award (R35GM138036 to C.A.D.) and R01HL085339 (to A.J.P.), and fellowship support for E.M. and J.E.V. through the NSF GRFP under Grant No. DGE 2241144. The authors gratefully acknowledge the following researchers at the University of Michigan: Jingjing Chen & Professor Jon Estrada for their assistance with high-speed imaging of droplet formation; Sungwan Park & Professor Albert T. Liu for their assistance with interfacial tension measurements, and Yiqun Li & Professor Abdon Pena-Francesch for their assistance with viscosity measurements. We also extend our sincere thanks to Professor Maria M. Coronel and her laboratory at the University of Michigan for the discussion on device workflow. Finally, we are grateful to Priyan Weerappuli for his insightful feedback and contributions in improving the efficiency of device fabrication. Figures were generated with GraphPad Prism, Python, and BioRender.

## References

[1] O. Shofolawe-Bakare, N. D. Leipzig, Acta Biomaterialia 2026, 213, 103.

[2] S. Feng, K. Chen, S. Wang, Adv Healthc Mater 2025, 14, e01947.

[3] A. Vaziri, R. Maia, P. Zhang, L. Agresti, J. Sjollema, M.-A. Shahbazi, H. A. Santos, Advanced Healthcare Materials 2026, 15, e02462.

[4] L. Riley, L. Schirmer, T. Segura, Current Opinion in Biotechnology 2019, 60, 1.

[5] S. Xin, D. Chimene, J. E. Garza, A. K. Gaharwar, D. L. Alge, Biomater. Sci. 2019, 7, 1179.

[6] A. J. Seymour, S. Shin, S. C. Heilshorn, Advanced Healthcare Materials 2021, 10, 2100644.

[7] I. W. Zhang, L. S. Choi, N. E. Friend, A. J. McCoy, F. S. Midekssa, M. M. Hu, E. Alsberg, S. C. Lesher-Pérez, J. P. Stegemann, B. M. Baker, A. J. Putnam, Acta Biomaterialia 2025, 201, 283.

[8] L. G. Brunel, F. Christakopoulos, D. Kilian, B. Cai, S. M. Hull, D. Myung, S. C. Heilshorn, Adv Healthc Mater 2024, 13, e2303325.

[9] D. J. Shiwarski, A. R. Hudson, J. W. Tashman, A. W. Feinberg, APL Bioeng 2021, 5, 010904.

[10] J. C. De La Vega, P. Elischer, T. Schneider, U. O. Häfeli, Nanomedicine 2013, 8, 265.

[11] A. Kalantarifard, E. Alizadeh-Haghighi, C. Elbuken, Chemical Engineering Science 2022, 261, 117947.

[12] A. C. Daly, L. Riley, T. Segura, J. A. Burdick, Nat Rev Mater 2020, 5, 20.

[13] O. Shofolawe-Bakare, N. D. Leipzig, Acta Biomaterialia 2026, 213, 103.

[14] A. Ofner, D. G. Moore, P. A. Rühs, P. Schwendimann, M. Eggersdorfer, E. Amstad, D. A. Weitz, A. R. Studart, Macromolecular Chemistry and Physics 2017, 218, 1600472.

[15] E. Stolovicki, R. Ziblat, D. A. Weitz, Lab Chip 2017, 18, 132.

[16] Z. Li, A. M. Leshansky, L. M. Pismen, P. Tabeling, Lab Chip 2015, 15, 1023.

[17] Z. Liu, C. Duan, S. Jiang, C. Zhu, Y. Ma, T. Fu, Journal of Industrial and Engineering Chemistry 2020, 92, 18.

[18] A. Montessori, M. Lauricella, E. Stolovicki, D. A. Weitz, S. Succi, Physics of Fluids 2019, 31, 021703.

[19] Z. Shi, X. Lai, C. Sun, X. Zhang, L. Zhang, Z. Pu, R. Wang, H. Yu, D. Li, 2020.

[20] A. Montessori, M. Lauricella, S. Succi, E. Stolovicki, D. Weitz, Phys. Rev. Fluids 2018, 3, 072202.

[21] M. L. Eggersdorfer, H. Seybold, A. Ofner, D. A. Weitz, A. R. Studart, Proceedings of the National Academy of Sciences 2018, 115, 9479.

[22] X. Xu, H. Yuan, R. Song, M. Yu, H. Y. Chung, Y. Hou, Y. Shang, H. Zhou, S. Yao, Biomicrofluidics 2018, 12, 014103.

[23] C. Roosa, L. Pruett, J. Trujillo, A. Rodriguez, B. Pfaff, N. Cornell, C. Flanagan, D. R. Griffin, Journal of Visualized Experiments (JoVE) 2022, e64119.

[24] J. M. de Rutte, J. Koh, D. Di Carlo, Advanced Functional Materials 2019, 29, 1900071.

[25] E. Amstad, M. Chemama, M. Eggersdorfer, L. R. Arriaga, M. P. Brenner, D. A. Weitz, Lab Chip 2016, 16, 4163.

[26] C. Roosa, L. Pruett, J. Trujillo, A. Rodriguez, B. Pfaff, N. Cornell, C. Flanagan, D. R. Griffin, Journal of Visualized Experiments (JoVE) 2022, e64119.

[27] G. T. Vladisavljevic, I. Kobayashi, M. Nakajima, Powder Technology 2008, 183, 37.

[28] M. Schulz, F. von Stetten, R. Zengerle, N. Paust, Langmuir 2019, 35, 9809.

[29] C. H. Y. Chung, B. Cui, R. Song, X. Liu, X. Xu, S. Yao, Micromachines 2019, 10, 592.

[30] H. Ji, J. Lee, J. Park, J. Kim, H. S. Kim, Y. Cho, Biosensors 2022, 12, 118.

[31] N. Raval, R. Maheshwari, D. Kalyane, S. R. Youngren-Ortiz, M. B. Chougule, R. K. Tekade, In Basic Fundamentals of Drug Delivery (Ed.: Tekade, R. K.), Academic Press, 2019, pp. 369–400.

[32] S. Barkley, S. J. Scarfe, E. R. Weeks, K. Dalnoki-Veress, Soft Matter 2016, 12, 7398.

[33] Z. Li, S. Y. Mak, A. Sauret, H. C. Shum, Lab Chip 2014, 14, 744.

[34] B. E. Campbell, K. Zhang, A. Shi, S. Rostami, D. Pioche-Lee, C. Li, A. Leblond, A. Forigua, C.-M. Boghdady, C. Moraes, S. C. Lesher-Pérez, ACS Appl. Bio Mater. 2025, 8, 3801.

[35] S. Utech, R. Prodanovic, A. S. Mao, R. Ostafe, D. J. Mooney, D. A. Weitz, Advanced Healthcare Materials 2015, 4, 1628.

[36] C. A. DeForest, D. A. Tirrell, Nature Mater 2015, 14, 523.

[37] R. Rizzo, D. Ruetsche, H. Liu, M. Zenobi-Wong, Advanced Materials 2021, 33, 2102900.

[38] E. M. Plaster, M. K. Eiken, C. Loebel, Carbohydrate Polymer Technologies and Applications 2023, 6, 100360.

