## Supplemental Figures Document for "A robust and user-agnostic step-emulsion platform for scalable microgel fabrication"

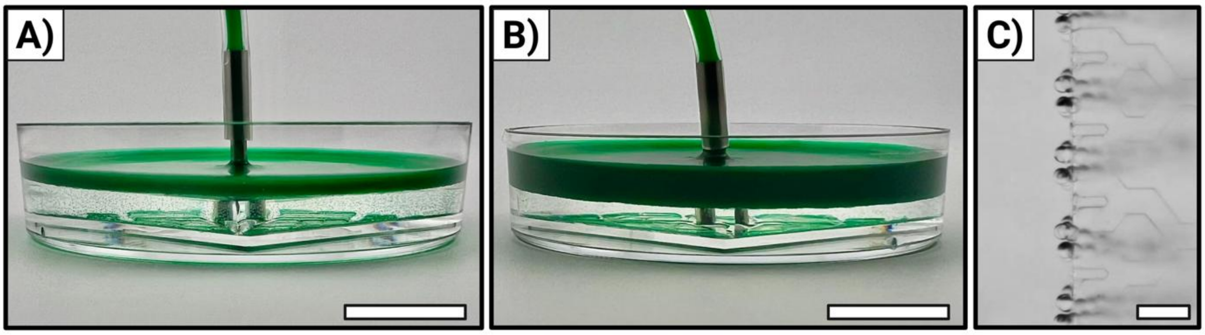


**Figure S1.** Images of droplet generation of water at 10 mL hr$^{-1}$ using a 30h150w device. A) Image after perfusing 2 mL of solution (t = 12 min); scale bar is 15 mm. B) Image after perfusing 5 mL of solution (t = 30 min); scale bar is 15 mm. C) Image of droplet generation at the outlet channel; scale bar is 300 µm. A and B use green food dye diluted in water for visualization and C uses only water.

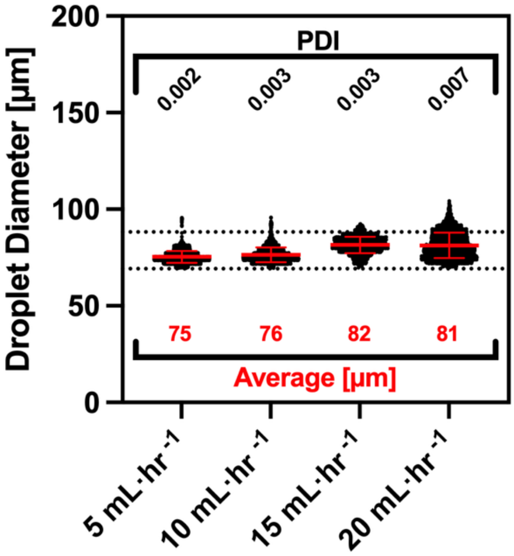


**Figure S2.** Scatter dot plots of the Gaussian distributions shown in Figure 1F (ordered in increasing flow rate). Droplet populations were produced from a 512-outlet channel device with outlet channel dimensions of 20 µm height and 120 µm width and all were infused with 10 wt% PEG-NB hydrogel precursor solution. Dashed lines indicate the tolerance interval representing 90% of the droplet population with 95% confidence, 69.2 to 88.2 µm. Polydispersity index (PDI) is reported above the scatter dot plots (in black text) and average droplet diameter (µm) below (in red text)

**Table S1.** Corresponding numerical data for Figure 2 listing channel dimensions, tested flow rates with corresponding flow velocities, and droplet size outcomes.

| Aspect Ratio Name | Height [μm] | Width [μm] | Aspect Ratio (h/w) | Tested Flow Rate [mL/hr] | Corresponding Flow Velocity [mm/sec] | Sample Size | Droplet Diameter [μm] | Droplet Diameter StDev [μm] | PDI |
|---|---|---|---|---|---|---|---|---|---|
| 1/12 | 15.0 | 176 | 0.09 | 0.5 | 1.6 | 3337 | 53.5 | 2.7 | 0.003 |
| | | | | 1 | 3.3 | 1436 | 52.8 | 2.9 | 0.006 |
| | | | | 1.5 | 4.9 | 1063 | 71.5 | 6.7 | 0.008 |
| | | | | 2 | 6.6 | 601 | 74.3 | 34.8 | 0.012 |
| | | | | 5 | 16.5 | 1825 | 72.2 | 40.2 | 0.032 |
| | | | | 10 | 32.9 | 2224 | 70.9 | 42.5 | 0.087 |
| 1/6 | 21.5 | 120 | 0.18 | 0.5 | 1.7 | 21181 | 70.6 | 5.0 | 0.005 |
| | | | | 1 | 3.4 | 9836 | 69.1 | 5.1 | 0.005 |
| | | | | 1.5 | 5.0 | 11917 | 71.0 | 8.2 | 0.013 |
| | | | | 2 | 6.7 | 4071 | 94.5 | 35.7 | 0.143 |
| | | | | 5 | 16.8 | 9954 | 211.8 | 176.9 | 0.697 |
| | | | | 10 | 33.7 | 9501 | 250.3 | 183.7 | 0.539 |
| 1/3 | 29.7 | 88 | 0.34 | 0.5 | 1.7 | 3055 | 91.4 | 2.1 | 0.001 |
| | | | | 0.75 | 2.5 | 2228 | 87.7 | 5.6 | 0.004 |
| | | | | 1 | 3.3 | 2127 | 92.3 | 6.1 | 0.004 |
| | | | | 2 | 6.7 | 345 | 166.2 | 45.6 | 0.075 |
| | | | | 3.5 | 11.6 | 160 | 237.8 | 95.5 | 0.161 |
| | | | | 5 | 16.6 | 2024 | 309.6 | 190.3 | 0.378 |
| | | | | 10 | 33.3 | 1738 | 279.5 | 205.2 | 0.540 |
| 1/2 | 36.6 | 71.8 | 0.51 | 0.5 | 1.7 | 911 | 123.5 | 3.8 | 0.001 |
| | | | | 1 | 3.3 | 1585 | 105.9 | 4.0 | 0.001 |
| | | | | 2 | 6.6 | 2627 | 101.8 | 10.5 | 0.011 |
| | | | | 5 | 16.5 | 439 | 145.9 | 61.9 | 0.180 |
| | | | | 10 | 33.1 | 1444 | 298.2 | 248.1 | 0.692 |
| 1/1 | 54.7 | 50.8 | 1.08 | 0.5 | 1.6 | 1282 | 167.0 | 8.0 | 0.002 |
| | | | | 1 | 3.1 | 1157 | 183.1 | 15.4 | 0.007 |
| | | | | 1.5 | 4.7 | 1589 | 139.5 | 20.6 | 0.022 |
| | | | | 3.5 | 10.9 | 121 | 255.7 | 52.1 | 0.041 |
| | | | | 10 | 31.3 | 944 | 322.0 | 234.8 | 0.532 |
| 2/1 | 90.4 | 38.7 | 2.34 | 0.5 | 1.2 | 1274 | 140.2 | 8.0 | 0.003 |
| | | | | 1 | 2.5 | 612 | 138.5 | 10.4 | 0.006 |
| | | | | 1.5 | 3.7 | 783 | 140.0 | 12.5 | 0.008 |
| | | | | 2 | 5.0 | 660 | 148.0 | 16.0 | 0.012 |
| | | | | 5 | 12.4 | 585 | 169.3 | 30.1 | 0.032 |
| | | | | 10 | 24.8 | 527 | 182.5 | 53.8 | 0.087 |

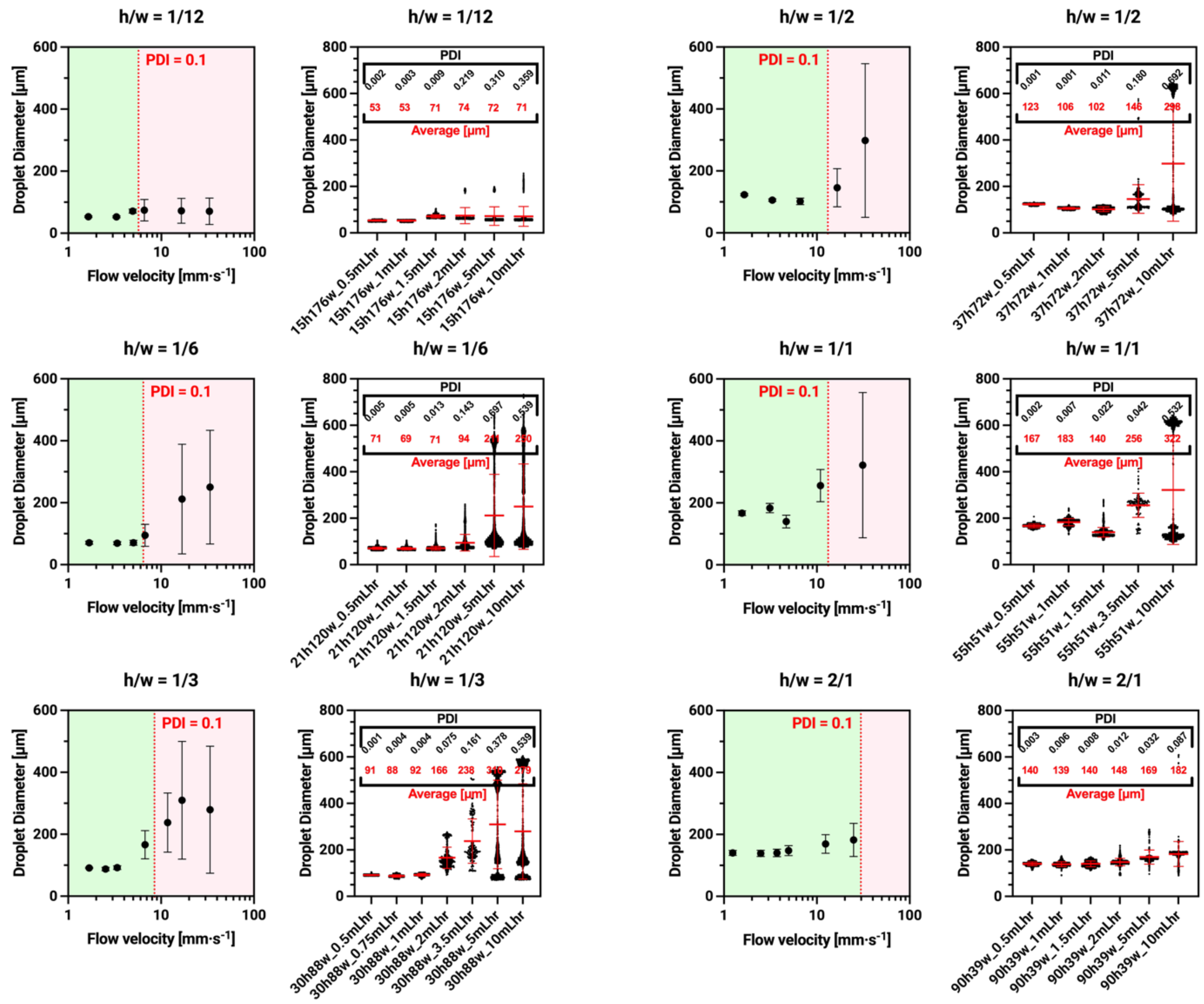

**Figure S3.** Resulting droplet diameter as a function of increasing flow rate and the same data plotted as scatter dot plots presenting droplet distribution. At lower aspect ratios, the device follows flow independence similar to literature. As aspect ratio increases, the transition into polydispersity is gradual rather than drastic as seen in lower aspect ratios. Droplet populations were produced from a 32-outlet channel device with varying outlet channel dimensions and all were infused with 10 wt% PEG-NB hydrogel precursor. Polydispersity index (PDI) is reported in black text and average droplet diameters (μm) in red text above the scatter dot plots.

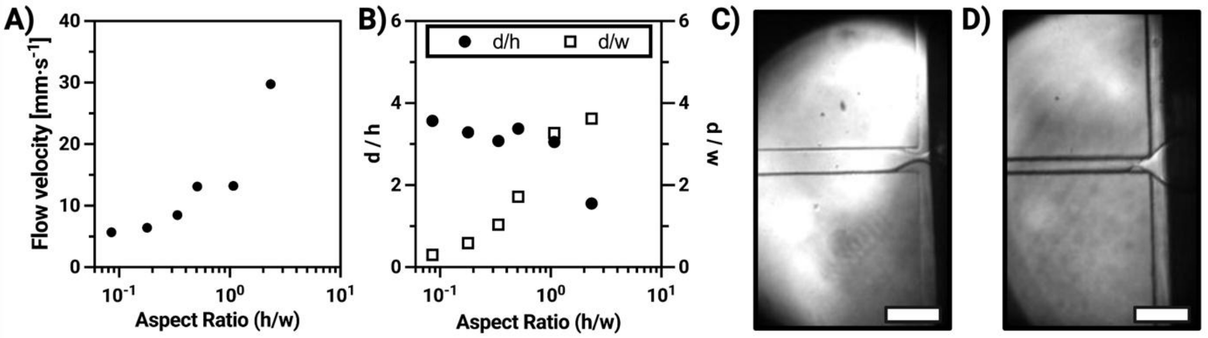


**Figure S4.** A) At a similar cross-sectional area, increases in aspect ratio increase the max monodisperse flow velocity. Here, transitional flow velocities are estimated from logistical regression and plotted against their respective aspect ratios. B) Ratio of monodisperse droplet diameter to either width or height plotted against their respective aspect ratios. Resulting droplet diameter $d$ is roughly 3 to 4 times the smallest dimension of the outlet channel (which is height for aspect ratios < 1 and width when aspect ratio > 1). C) High-speed imaging of droplet pinching for 1/2 (h/w) aspect ratio. D) High-speed imaging of droplet pinching for 2/1 (h/w) aspect ratio; scale bars are 150 µm. A, B, C, and D all used 10 wt% PEG-NB hydrogel precursor solution.

**Table S2.** Predicted transitional flow velocities with the fitted logistic regression parameters for each model shown corresponding to Figure 2.

| Aspect Ratio Name | Height [µm] | Width [µm] | Aspect Ratio (h/w) | Logistical Regression Equation | s | k | $x_0$ | c | Predicted Transitional Velocity [mm/sec] |
|---|---|---|---|---|---|---|---|---|---|
| 1/12 | 15.0 | 176 | 0.09 | $log_{10}y = s \cdot \arctan(k \cdot x + x_0) + c$ | -0.70 | -2.53 | 13.35 | -1.56 | 5.7 |
| 1/6 | 21.5 | 120 | 0.18 | $log_{10}y = s \cdot \arctan(k \cdot x + x_0) + c$ | -0.74 | -1.04 | 6.21 | -1.33 | 6.4 |
| 1/3 | 29.7 | 88 | 0.34 | $log_{10}y = s \cdot \arctan(k \cdot x + x_0) + c$ | 236.03 | 21.40 | 34.50 | -370.7 | 8.5 |
| 1/2 | 36.6 | 71.8 | 0.51 | $log_{10}y = s \cdot \arctan(k \cdot x + x_0) + c$ | -1.68 | -0.13 | 0.67 | -2.37 | 13.1 |
| 1/1 | 54.7 | 50.8 | 1.08 | $log_{10}y = s \cdot \arctan(k \cdot x + x_0) + c$ | 418.98 | 12.00 | 118.40 | -657.6 | 13.2 |
| 2/1 | 90.4 | 38.7 | 2.34 | $log_{10}y = s \cdot \arctan(k \cdot x + x_0) + c$ | 252.15 | 10.75 | 109.25 | -396.5 | 29.8 |

**Table S3.** Key for how the geometric parameters were calculated.

| Geometric Parameter | Unit | Variable | Equation |
|---|---|---|---|
| Channel Height | µm | $h$ | - |
| Channel Width | µm | $w$ | - |
| Cross-Sectional Area | $µm^2$ | $CSA$ | $CSA = h \cdot w$ |
| Aspect Ratio (h/w) | - | $AR$ | $AR = \frac{h}{w}$ |
| Normalized Aspect Ratio | - | $AR_n$ | $AR_n = \begin{cases} \frac{h}{w}, & if\ h > w \\ \frac{-h}{w}, & if\ h < w \end{cases}$ |
| Perimeter | µm | $l_{peri}$ | $l_{peri} = 2h + 2w$ |
| Diagonal | µm | $l_{diag}$ | $l_{diag} = \sqrt{h^2 + w^2}$ |
| Smallest Dimension | µm | $l_{min}$ | $l_{min} = \begin{cases} w, & if\ h > w \\ h, & if\ h < w \end{cases}$ |
| Largest Dimension | µm | $l_{max}$ | $l_{max} = \begin{cases} h, & if\ h > w \\ w, & if\ h < w \end{cases}$ |
| Radius of Cross-Sectional Area | µm | $r_{CSA}$ | $r_{CSA} = \sqrt{\frac{CSA}{\pi}}$ |
| Average Dimension | µm | $l_{avg}$ | $l_{avg} = \frac{h + w}{2}$ |

| | | | |
|---|---|---|---|
| Hydraulic Radius | µm | $r_{hyd}$ | $r_{hyd} = \frac{h \cdot w}{2(h+w)}$ |

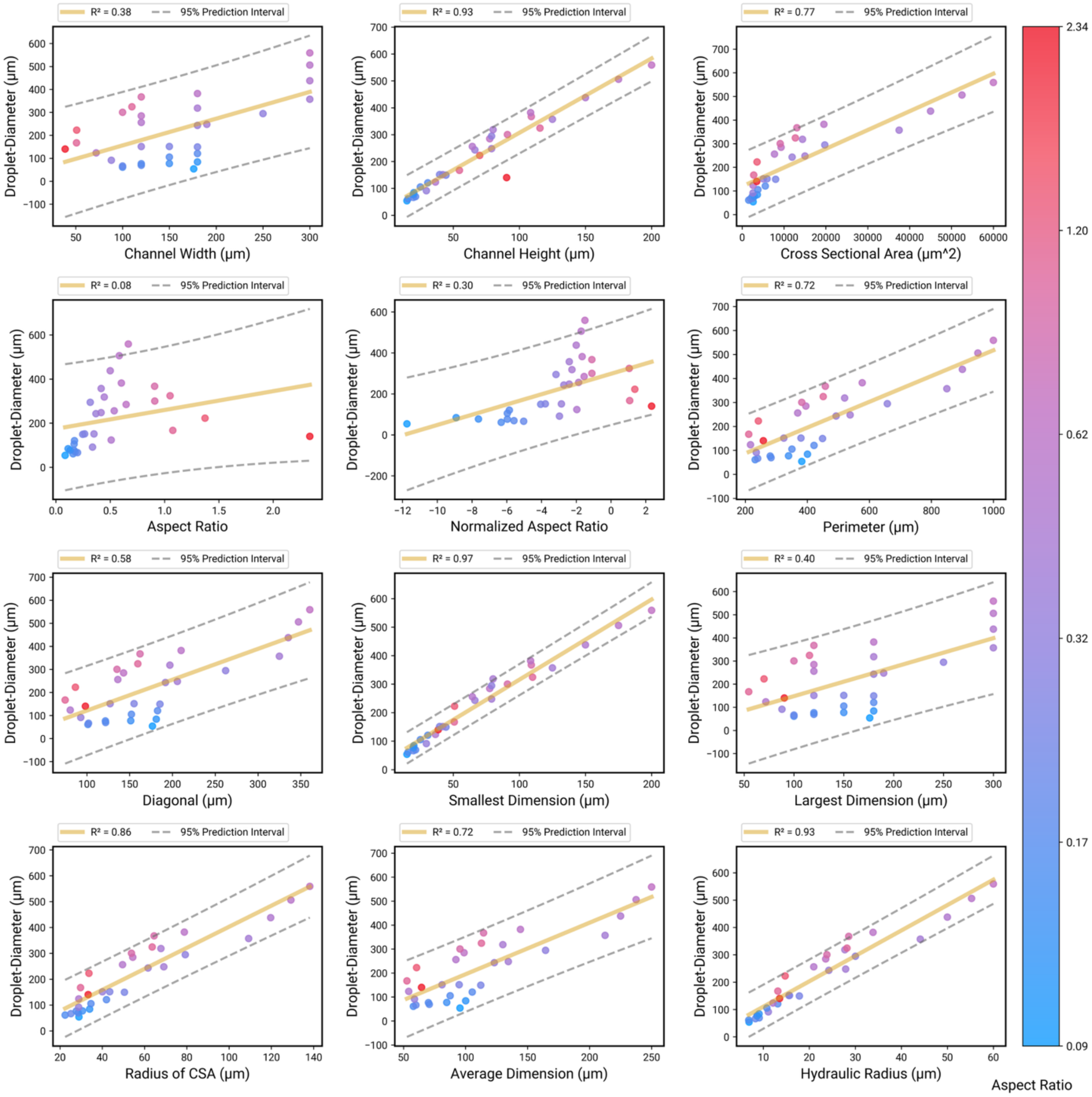


**Figure S5.** Various geometric parameters were tested for a correlation to droplet size with smallest dimension demonstrating the strongest correlation.

**Table S4.** Key for materials tested and the corresponding figures and tables the materials were used in.

| Material Tested | Formulation | Shorthand | Gelation Chemistry | Figure and Table Ref. |
|---|---|---|---|---|
| Water | - | Water | - | Figures 1B, 5A, 6A, 6B, and S1<br>Table S5 |
| Phosphate buffered saline | - | PBS | - | Figure 5A |
| 8-arm Poly(ethylene glycol) norbornene | 10 wt% | PEG-NB | Thiol–ene photopolymerization (step-growth) | Figures 1C, 1E, 1F, 2B, 2C, 3A, 3B, 3C, 4A, 4B, 5A, 5B, 6A, 6B, S2, S3, S4A, S4B, S4C, S4D, and, S5<br>Tables S1, S2, and S5 |
| Trehalose | 1 M | Trehalose | - | Figure 5A |
| Alginate | 2, 3, 4, and 5 wt% | Alginate | Ionic crosslinking | Figures 1D, 5A, 6A, and 6B<br>Table S5 |
| Gelatin-methacrylate | 5 % (w/v) | GelMA | Free-radical photopolymerization | Figures 1D and 5A |
| Poly(ethylene glycol) tetrabicyclononyne | 4 mM | PEG-BCN | Strain-Promoted Azide-Alkyne Cycloaddition (SPAAC) | Figures 1D and 5A |
| Gelatin-norbornene | 10 % (w/v) | GelNB | Thiol–ene photopolymerization (step-growth) | Figure 5A |
| Poly(ethylene glycol) diacrylate | 10 % (w/v) | PEGDA | Free-radical photopolymerization | Figure 5A |
| Norbornene-modified hyaluronic acid | 2 % (w/v) | NorHA | Thiol–ene photopolymerization (step-growth) | Figures 1D and 5A |
| 8-arm Poly(ethylene glycol) maleimide | 10 wt% | PEG-MAL | Thiol–maleimide Michael-type addition | Figure 5A |
| 8-arm Poly(ethylene glycol) vinyl sulfone | 5 wt% | PEG-VS | Thiol–vinyl sulfone Michael-type addition | Figure 5A |

**Table S5.** The droplet generator is compatible with more viscous materials, and as long as the user works below a critical capillary number, monodisperse droplets are achievable. Fluid properties of various materials and tested flow rates based on an estimated critical capillary number (Ca = 0.014) from literature[20] are reported here.

| Material | Concentration | Viscosity [mPa s] at ~25 °C | Interfacial Tension [mN/m] | Estimated Flow Rate [mL/hr] | Flow Rate Tested [mL/hr] | Droplet Diameter [μm] | PDI |
|---|---|---|---|---|---|---|---|
| Water | - | 1 | 5.0 | 265 | 10 | 79 | 0.003 |
| PEG-NB | 10 wt% | 9 | 2.9 | 20 | 10 | 76 | 0.003 |
| Alginate | 2 wt% | 44 | 2.6 | 3.7 | 4 | 71 | 0.001 |
| Alginate | 3 wt% | 171 | 4.0 | 1.5 | 2 | 70 | 0.001 |
| Alginate | 4 wt% | 396 | 3.6 | 0.6 | 0.5 | 72 | 0.001 |
| Alginate | 5 wt% | 669 | 3.8 | 0.4 | 0.3 | 69 | 0.014 |
