## Supplementary material for "A robust and user-agnostic step-emulsion platform for scalable microgel fabrication": CAD Files, PDF of Device Design, Text Description: Pioche-Lee, et al_BDDG-CAD Files Description.txt

The folder contains two CAD files containing both a 3D printable 32-channel high-throughput device and an AutoCAD drawing file of a 512-channel device. Due to mold height resolution limitations, smaller outlet channel designs are made using SU8 photolithography and larger outlet channel dimensions (>100 µm height, >250 µm width) are possible with 3D printing. The AutoCAD file contains two photomask designs for a first and second layer. The second layer design is necessary for a defined cutting region and provides a uniform edge and outlet for droplet production. Additionally a PDF file is included as a representation of the two photomask layers. The following file names are listed below with a brief description. 


260430_LesherPerezLab_2LayerPhotomasks_DPL-Layout.pdf
A PDF of the two mask layers of the device, to visualize the device channel and layout. This design is specifically for a 150 μm channel width, for a 512 outlet channel device. 

260430_LesherPerezLab_2LayerPhotomasks_DPL.dwg
A .dwg file of the two mask layers of the device. This design is specifically for a 150 μm channel width, for a 512 outlet channel device. We have made this accessible in case others would like to do clean room fabrication of the system. 

260430_LesherPerezLab_3DP_DropletGenerator_DPL.stl
A 3D model of four, 32 channel droplet generators of different sizes (125 μm height x 300 μm width; 150 μm height x 300 μm width; 175 μm height x 300 μm width; 200 μm height x 300 μm width)
