## Supplementary figures and images for "A robust and user-agnostic step-emulsion platform for scalable microgel fabrication"

### 260430_LesherPerezLab_2LayerPhotomasks_DPL-Layout.pdf

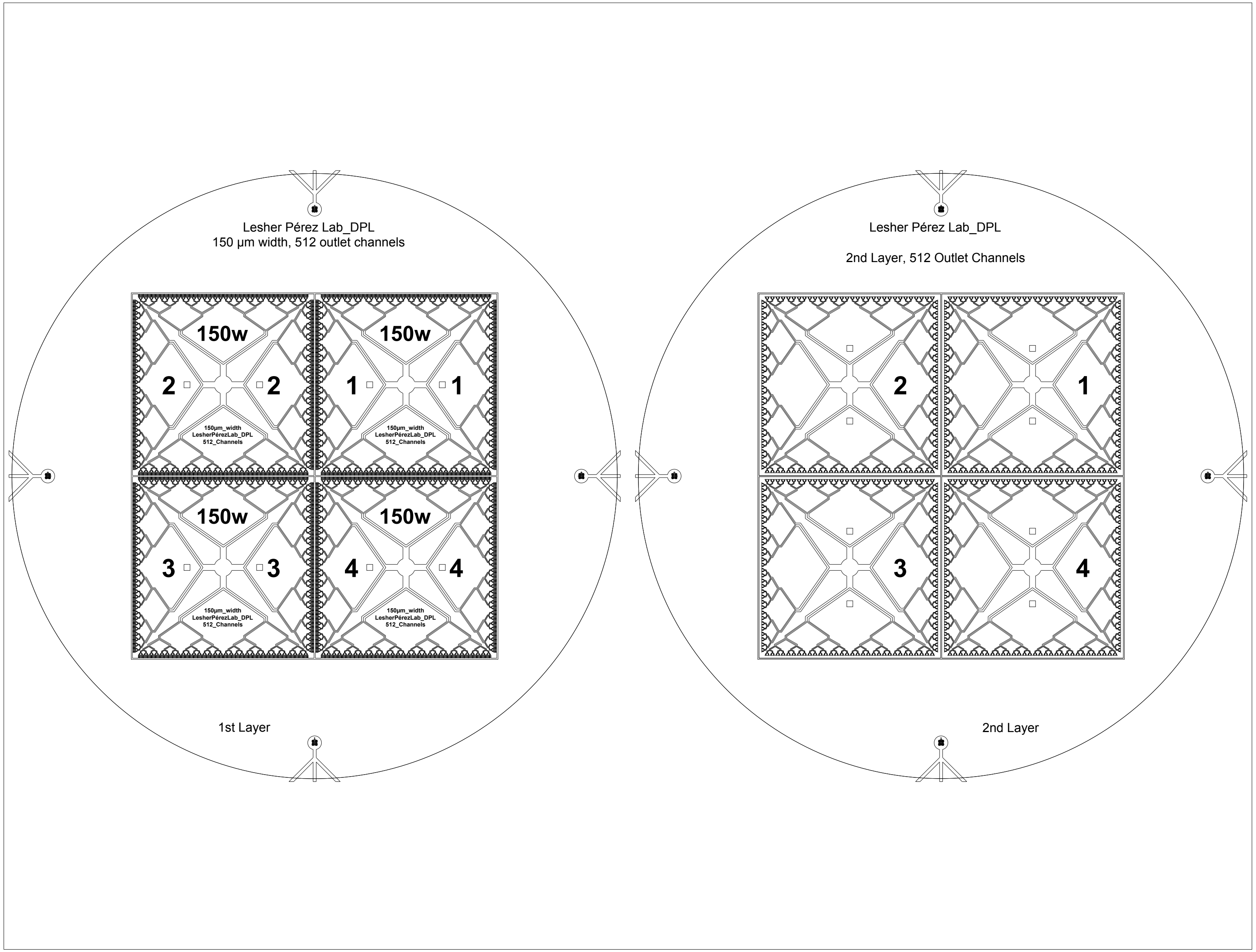
