## Supplementary material for "A robust and user-agnostic step-emulsion platform for scalable microgel fabrication": Droplet Generator Operating Protocol Provided to New Users

Date: 03/31/2026 | Last Updated: 04/12/2026
Written by: Durante Pioche-Lee; Reviewed by: Yoke Qi Ho, Wahidur Rahman, Sasha Cai Lesher-Pérez

### Simple Droplet Generation

#### Operating Procedure

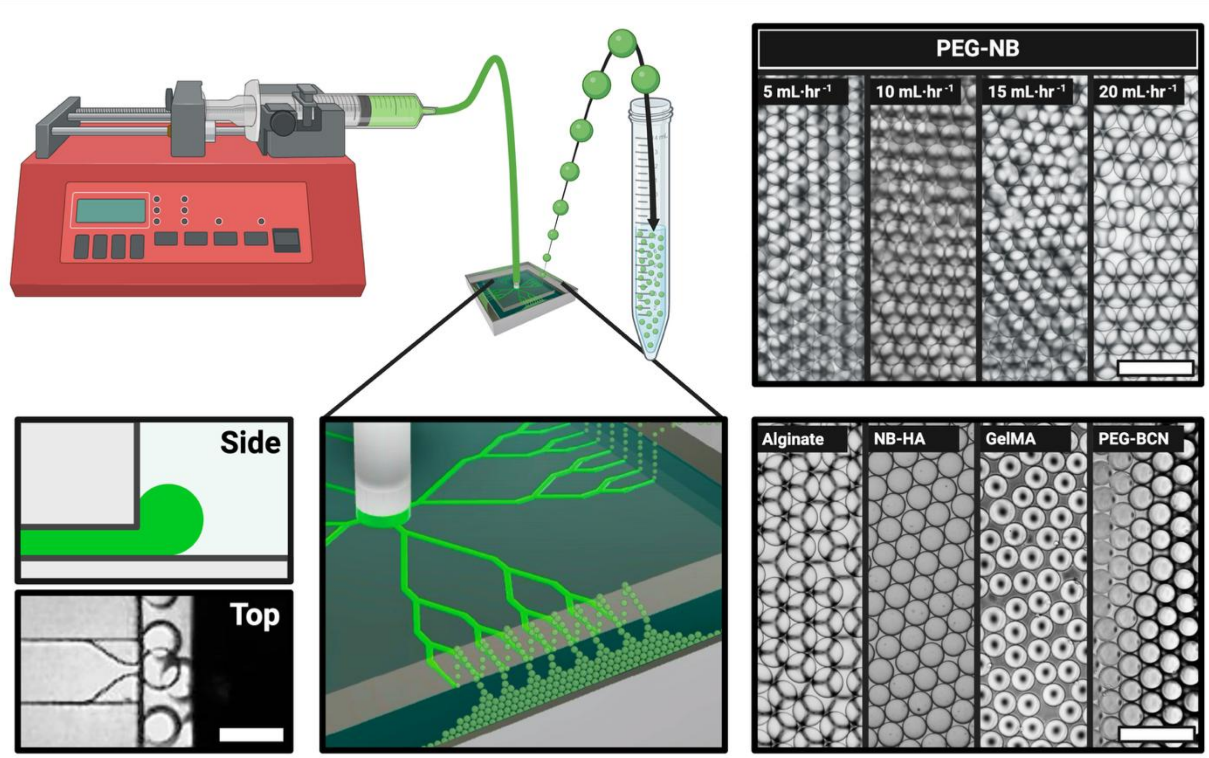


***Figure 1.*** *General overview of the single-input droplet generator. Precursor solution is supplied laterally and the produced droplets float to the top where collection and gelation happen. Monodisperse PEG-NB droplets were produced by infusion at various flow rates and using various hydrogel precursor solutions. Scale bars = 120 µm (bottom-left), 250 µm (top- and bottom-right).*

#### Description

This approach leverages the step-emulsion technology with two phases (gel precursor/dispersed phase and oil/continuous phase) to generate gel precursor droplets which detach from the bulk liquid and float to the surface. It utilizes ledge emulsification channels with a single syringe pump supplying active flow to parallelized outlet channels (

Figure **1**). Gel precursor is supplied to the polydimethylsiloxane (PDMS) device positioned in a static reservoir of high density oil with surfactant. Generated droplets can then undergo size characterization and gelation. Devices can be ordered here, through academic licensing of a core service by the Lesher- Pérez Research Group.

#### Equipment & Materials

- Optically clear 60mm diameter petri dish
- Packing tape
- Double-sided tape (Scotch tape)
- Vacuum desiccator
- Luer-Lock syringes (1 mL, 3 mL, etc.,)
- 14 gauge Luer-Lock blunt tips (link here)
- 50mL conical tube
- Dispersed phase: Gel Precursor
- *Optional: 50mL conical tube size strainer
- *Optional: 5 to 7” piece of tubing with an ID of 1.6mm (0.0625”) (link here)
- Static reservoir/continuous phase: high density oil with surfactant (Ex. 0.4% (w/w)

- 14 gauge cannula (attainable from the blunt tip above)
- PDMS droplet generation device plasma bonded to a PDMS coated glass slide (vacuum desiccate upon device arrival).

Pico-Surf® (PS) in Novec™ 7500 or 2% (w/w) RAN 008-FluoroSurfactant in HFE 7500)

#### PPE & Safety Considerations

**Caution:** Device bottom is cut glass and therefore poses a **SHARPS HAZARD.** If the dispersed phase clogs the channels, the system may pressurize and expel the dispersed phase. Removal of the cannula from the device while there is a positive pressure will also expel dispersed phase. Lab coats, goggles, gloves should be worn when working with the system.

#### Device Specification

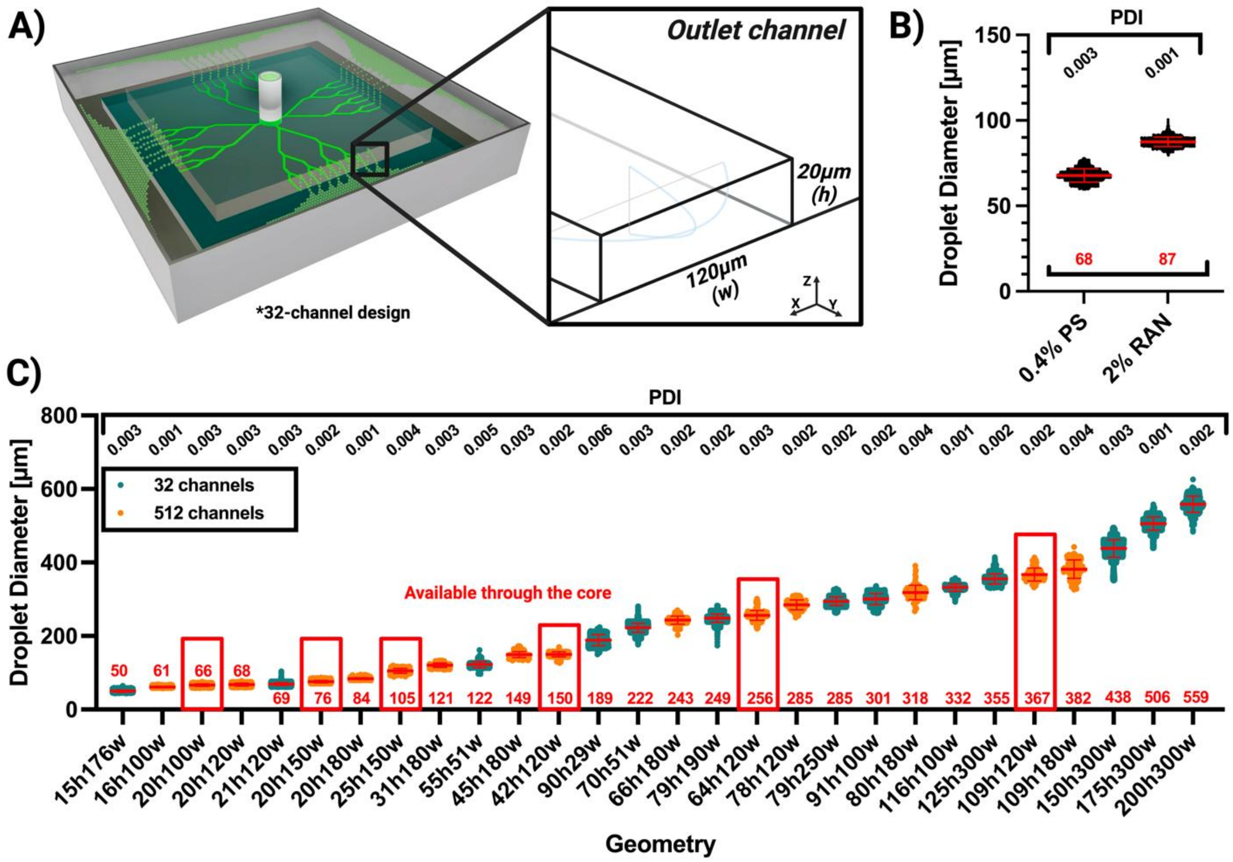


*Figure 2.*
*(A) Schematic of droplet generation highlighting outlet geometry with outlet channel: 20 µm (height) x 120 µm (width). (B) Surfactant type impacts droplet size.* 10wt% PEG-8-Norbornene at a 0.4 R-ratio was infused at 10 mL/hr using a device with 512-outlet channels of *20 µm height and 120 µm width device. Continuous phase solutions are 0.4% (w/w) Pico-Surf® (PS) in Novec™ 7500 and 2% (w/w) RAN 008-FluoroSurfactant in Novec™ 7500. (C)* Resulting droplet populations using various outlet geometries. Polydispersity index (PDI) is reported above in black text and average diameter is reported below in red text. Devices outlined in red boxes indicate devices available from the core. 512-channel data is considered high-throughput (capable of at least 10 mL/hr infusion rates). All devices were characterized with 10wt% PEG-8-Norborene at a 0.4 r-ratio using a 0.4% (w/w) Pico-Surf® in Novec™ 7500 as the continuous phase.

### Procedure

*This procedure is intended as general guidance for users. While there are variations of this procedure (e.g. manual injection into the devices), this protocol describes our standard approach using a syringe pump to drive fluid flow into the droplet-generation device. The steps below provide a generalized method we found to work reliably.*

#### 1. Prepare the droplet generation device

##### 1.1 Assemble and degas the device

a. Remove any lint or debris from the PDMS device surface & PDMS coated glass slide with packing tape. Any remaining debris will be suspended in the oil.
b. Adhere a piece of double-sided tape to the bottom of the glass slide and firmly press the device (tape-side down) into the petri dish.
c. Let the assembly sit in the vacuum desiccator for at least 15 minutes before use.

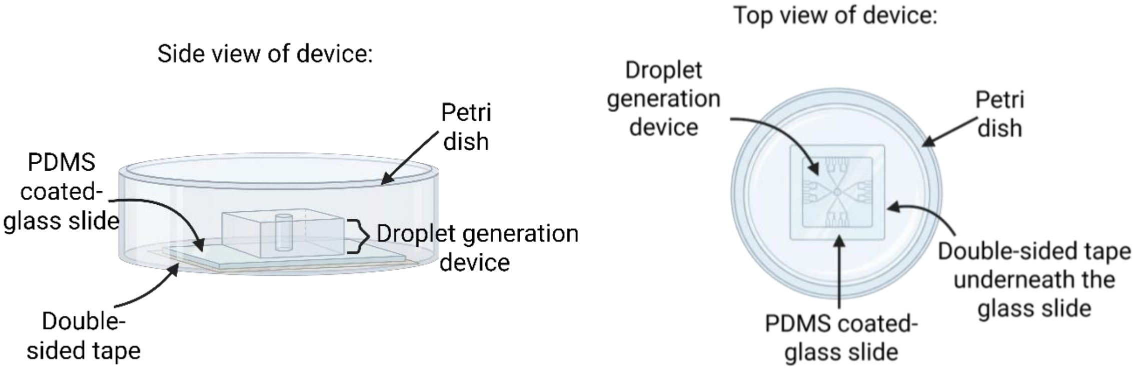


***Figure 3.*** *Images of the droplet generator's assembly from the side (left) and top (right) view. Created with Biorender.*

##### 1.2 Prepare the tubing

The device is supplied with an inlet punch made using a 2 mm biopsy punch. A 14G blunt tip provides snug connection into this inlet.

a. Remove a cannula from the blunt needle tip.
   i. Notes: Soaking the blunt tip in isopropyl alcohol overnight helps with removal.
   ii. For reference, 14G blunt tips match the [1/16” inner diameter tubing].
b. Place a blunt tip all the way into one end of the tubing and place the cannula tip (the metal tip) halfway into the other end of the tubing (as shown in Figure 4).
c. *User may also choose to directly inject gel precursor straight from the syringe blunt tip to the device. Instructions are provided in the next section.
   i. Pushing slow and controlled produces more monodisperse populations.

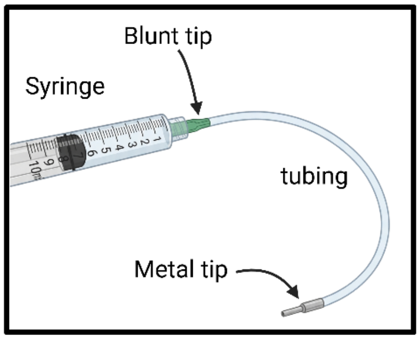


*Figure 4. The inlet (blunt tip) will be connected to the syringe, while the outlet (metal tip) will be pushed into the device.*

ii. For reference, pushing 50 µL every 10 seconds at a controlled infusion can be used for all devices except the 20h100w device which should be infused at 50 µL every 20 seconds.

#### 2. Setting up droplet generation with hydrogel precursor:

##### 2.1 Preparing the syringe

a. Spin down the gel precursor to remove any air bubbles.
b. Fill a syringe with gel precursor, making sure to minimize making small air bubbles.
    i. Use a syringe size that matches the infusion volume (e.g., for 800 µL of gel precursor, use a 1 mL syringe, not anything larger than a 1 mL syringe) to improve control over infusion due to fluctuations displacing smaller volumes with smaller syringes.
b. Hold the syringe upright (outlet up) and tap the syringe to break micro air bubbles.
    i. For manual injection, leave 100-200 µL volume of air above the gel precursor in the syringe. This air volume serves two purposes:
        i. The added capacitance will help dampen any fluctuations during manual infusion and eliminate dead volume. However, too large of an air section will increase the resistance and make manual infusion difficult.
c. For syringe pump-driven flow, attach the blunt tip with tubing to the syringe. Press the plunger until precursor is visible in the tubing. Ensure there are no air bubbles in the tubing.
d. Set up the syringe on the syringe pump. Ensure the correct syringe diameter is used (list of common syringe diameters here). Set the infusion rate.
e. Prime the syringe by operating the pump at a higher flow rate until the gel fills the rest of the tubing and outlet cannula. Pause until ready to start.

##### 2.2 Running the device

a. Remove the device assembly from the vacuum desiccator.
b. Add the 0.4% (w/w) Pico-Surf® (PS) in Novec™ 7500 or 2% (w/w) RAN 008-FluoroSurfactant in HFE 7500 into the petri dish until the liquid covers 1-2mm above the device (about 8-10 mL).
c. Ensure the oil phase coats the inner channels of the device by actively pipetting oil/continuous phase into the inlet.
    i. The previously performed vacuum desiccation will help remove any smaller residual air bubbles without needing to actively pipette the oil phase through the device.
d. Insert the cannula into the device inlet, but ensure the cannula is **not** pressed against the bottom (~0.5mm gap).
e. Begin infusion of the dispersed phase.
f. For syringe pump injection with tubing, once the syringe plunger reaches the end, reclaim the dispersed phase dead volume by:
    i. Ensure the syringe and tubing is maintained above the device throughout step f.
    ii. Separate the syringe from the blunt-tip assembly, while keep the tubing/blunt-tip above the device assembly.
    iii. Draw air into the syringe and reconnect the blunt-tip to the syringe.
    iv. Infuse the air to clear the dead volume of the dispersed phase.
    v. Stop when air is visibly pushed through the outlet channel.

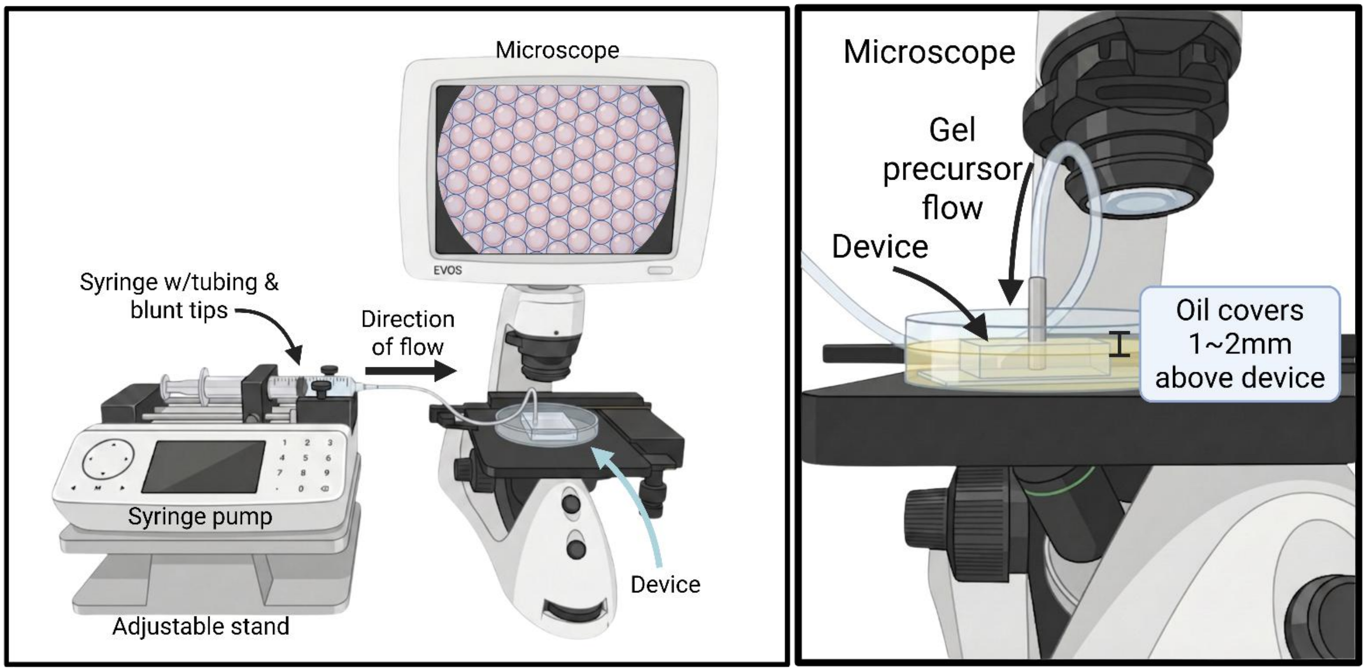


***Figure 5.*** *Example set-up for droplet generation. The device is set up to allow the gel precursor to flow into the device. The oil should cover the top of the device by 1~2mm. This ensures there is enough space for the droplets to float up.*

#### 3. Post droplet generation

a. Once 300-400 µL of gel precursor is infused, take droplet images with an appropriate objective (ensure droplets are not too small or big for the field of view) for image analysis of droplet diameter later. Ensure droplets are in the primary focal plane for size analysis (shown in Figure 6 & 11).
    i. 5 images is generally enough to reduce overlapping counts of droplets. Analyzing >100 droplets would be sufficient to represent the droplet diameter distribution.
b. Crosslink the droplet particles once droplet generation is completed or gel precursor is used up (shown in Figure 7). A list of gelation chemistries and their corresponding methods is described in our paper.
c. To recover microgels, attach a 1" 20G blunt tip to a 10 mL syringe and insert the blunt tip into the bottom the petri dish. Suck up most of the oil phase until less than 1 mL remains.
d. Once complete, pour the droplets from the petri dish into a 50mL conical tube. The oil phase will cling to the outside of the petri dish during pouring, to ensure minimal oil phase loss, carefully touch the petri dish bottom to the top of the conical tube, and slowly pour.
e. To reclaim residual droplets in the petri dish, use the reclaimed oil phase in the syringe to wash off remaining gels.
    i. Any unintended large droplets made can be filtered by pouring into a conical tube with a strainer.

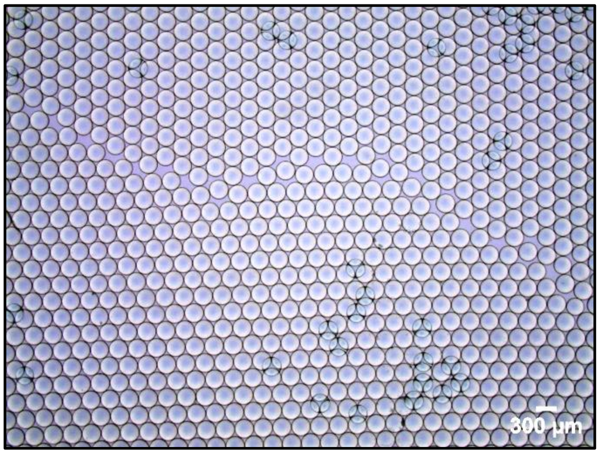


***Figure 6.*** *This is an image of droplets generated at 5mL/hr using an outlet channel with dimensions of 79 µm (h) x 190 µm (w). The gel precursor used is 10 wt% polyethylene glycol 8-arm norbornene (PEG8NB) - macromer, polyethylene glycol dithiol (PEGDT) - crosslinker, and lithium phenyl-2,4,6-trimethylbenzoylphosphinate (LAP) - photoinitiator.*

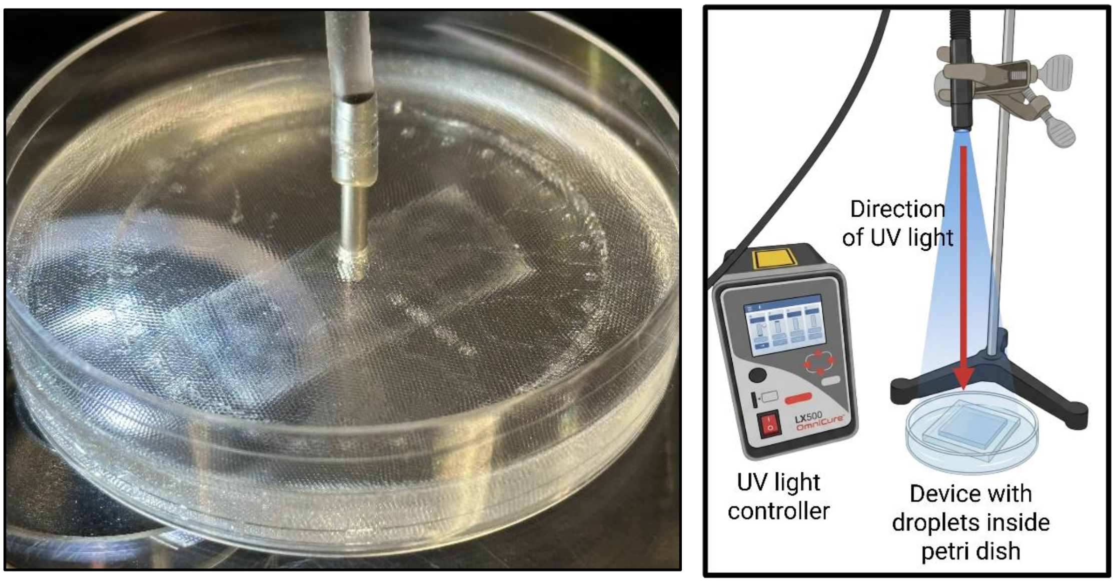


***Figure 7.*** *At the end of droplet generation, the droplets float to the top of the device (air/liquid) interface (left). UV light set-up to crosslink droplets into microgels (right).*

4. Analyzing droplet size generated:

<u>4.1 Setting up MATLAB script</u>

a. Using ImageJ, determine the pixel diameter length of the largest and smallest droplets. Also, put all images in one folder labeled "**Images**" (this example uses the name 'Images' seen in Fig. 10).

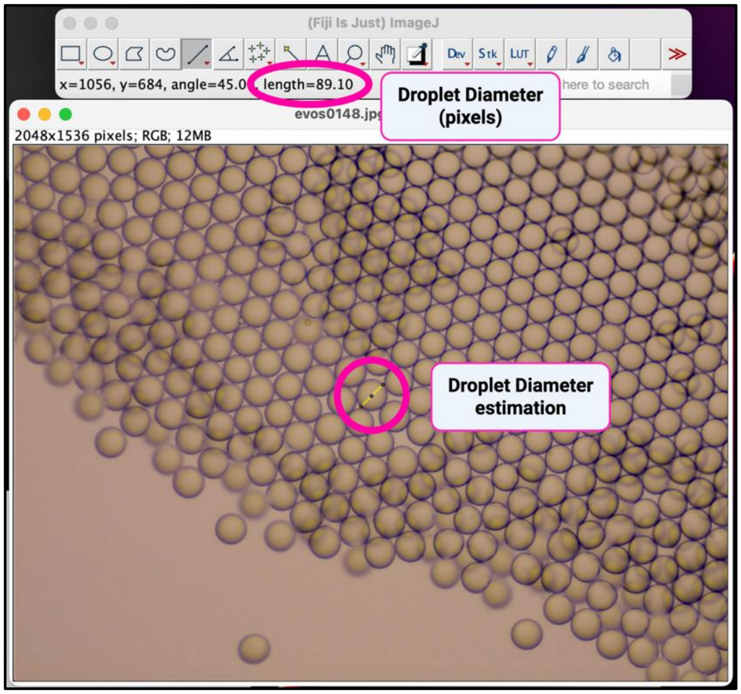


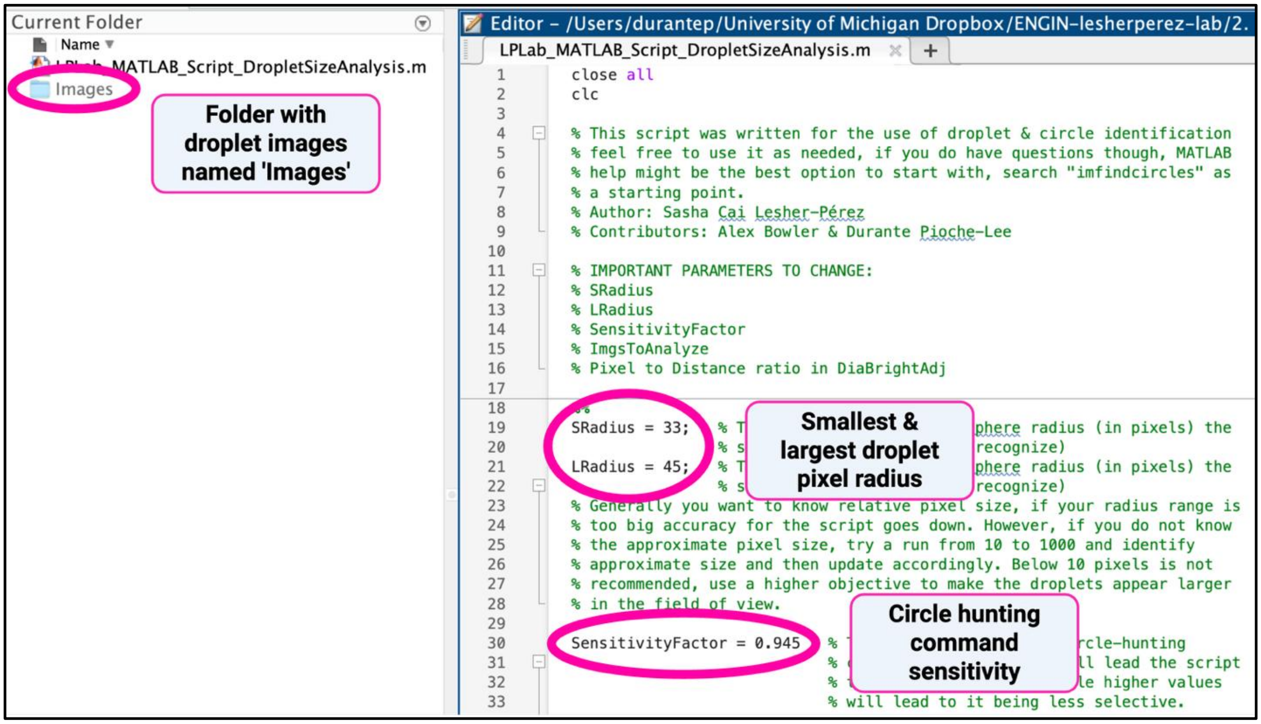


*Figure 8.* *Using ImageJ (or other photoviewer) to trace the pixel length and diameter of droplets.*

***Figure 9.*** *MATLAB script to track and analyze droplet diameter through detection and measurement of circle objects.*

b. In the script, input the largest and smallest pixel radius of droplets from the previous ImageJ analysis (Figure 10).
c. Input the sensitivity factor of the circle hunting command.
    i. **Note: the usual values used are between the range of 0.8~0.96. Close to 1 means more sensitive.
d. Ensure the file name that has the droplet images is named accordingly.
    i. *Note: the MATLAB script must be stored in the same directory as the droplet images ("Current Folder" top left).
e. Input the objective's pixel-to-distance conversion in line 173 (search "DiaBrightAdj").
f. Run the MATLAB script while the directory is open as seen in Figure 10 ("Current Folder" top left).
    i. **Note: if "imread error" pops up, change "for c = 3" value from 3 to 4 in line 95, also delete any "metadata" from the folder, it can't read it and it causes an error.

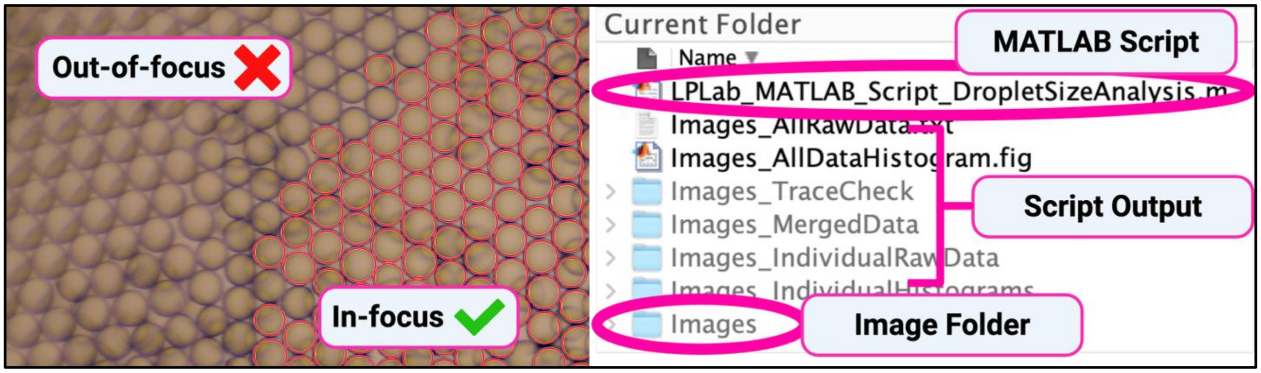


***Figure 10.*** *Circle function performed by the MATLAB code on the droplet images (left). Files generated after code is ran (right).*

#### 4.2 Analysis of average droplet diameter, PDI and CV.

a. Using excel, import the raw data of droplet diameter.
b. To calculate Polydispersity index (PDI) and coefficient of variance (CV), use:

$$PDI = \left(\frac{standard\ deviation\ of\ droplet\ diameter}{avg\ droplet\ diameter}\right)^2$$

$$CV = \frac{standard\ deviation\ of\ droplet\ diameter}{avg\ droplet\ diameter} * 100\%$$

c. To calculate the average droplet diameter, average out the raw data from one or multiple runs. Also calculate the standard deviation using the same data set.
d. Transfer data into GraphPad Prism or a similar data analysis software.
