## Supplementary material for "A robust and user-agnostic step-emulsion platform for scalable microgel fabrication": Microgel Purification Protocol Provided to New Users

Created 02/09/2023
Author: Durante Pioche-Lee
Lesher-Pérez Lab

### Microgel Purification

**Standard Operating Procedure**

#### Description

This protocol outlines two methods for purifying microgels after their collection in a dense oil phase (particularly 0.4wt% Pico-Surf® in Novec™ 7500). The first method is optimized for size exclusion or sorting which is ideal for applications requiring distinct microgel sizes, though it generally requires more time to complete. The second method bypasses size exclusion and is suitable when dealing with uniform populations, allowing all microgels to be used directly without additional sorting. Both methods assume the microgels have fully gelled but remain unhydrated (i.e., no swelling).

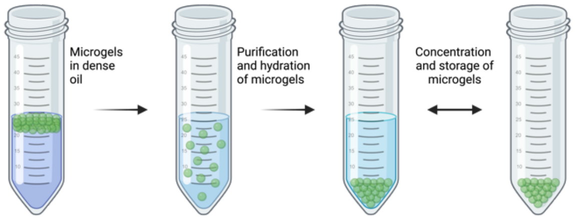


***Figure 1.*** *Overview of microgel purification*

#### Equipment & Materials

Below is a list of equipment and materials common to both methods. Method-specific materials are noted beside the item. Ensure material compatibility (especially for Hexane).

- Pure Novec™ 7500
- 1H, 1H, 2H, 2H-Perfluorooctanol (PFO) (link here)
- Hexanes (link here)
- 50mL conical tubes
- Pipettes and pipette tips
- Centrifuge for 50mL conical tubes
- Buffer (PBS, HEPES, etc.)
- Positive displacement pipettes and tips
- 0.45µm syringe filters
- Luer-Lock syringes
- 50mL conical tube-compatible mesh strainers (pluriStrainer link here) ***For method 1 only
- 22 gauge Luer-Lock blunt tips with 1” or 1.5” long tips ***For method 2 only

#### PPE & Safety Considerations

Lab coats, goggles, and appropriate gloves should be worn when purifying the microgels. With enough force, 22G blunt tips may puncture skin and therefore poses a **SHARPS HAZARD**. Additionally, centrifugation at high speeds poses a **PHYSICAL HAZARD;** ensure proper maintenance and operating protocols are used. *****IMPORTANT**: Proper university/laboratory-specific documentation and procedures should be written and followed when working with Hexane. Hexane is toxic, highly flammable, and its vapors can be explosive. Hexane causes skin and eye irritation and may cause drowsiness or dizziness. Hexane is suspected of damaging fertility or the unborn child. Hexane may be fatal if swallowed and enters airways. Hexane may cause damage to organs through prolonged or repeated exposure. All work done with hexane should be completed in a certified fume hood with safety glasses, flame-resistant

Created 02/09/2023
Author: Durante Pioche-Lee
Lesher-Pérez Lab
lab coat, and nitrile gloves (chemical resistant if possible like Microflex 93-260 [I-280], N-Dex Plus 8005 [I-85], TouchNTuff 92-600/650 [1-480], inspect gloves before use). Use Hexane-compatible materials when handling. Avoid inhalation and contact when working with Hexane. When done working with Hexane, store properly, dispose of waste following university hazardous waste procedures, and wash hands and forearms thoroughly.[1] **Again, follow laboratory-specific safety protocols and documentation when working with hexane.**

### Procedure – Method 1 (Size exclusion)

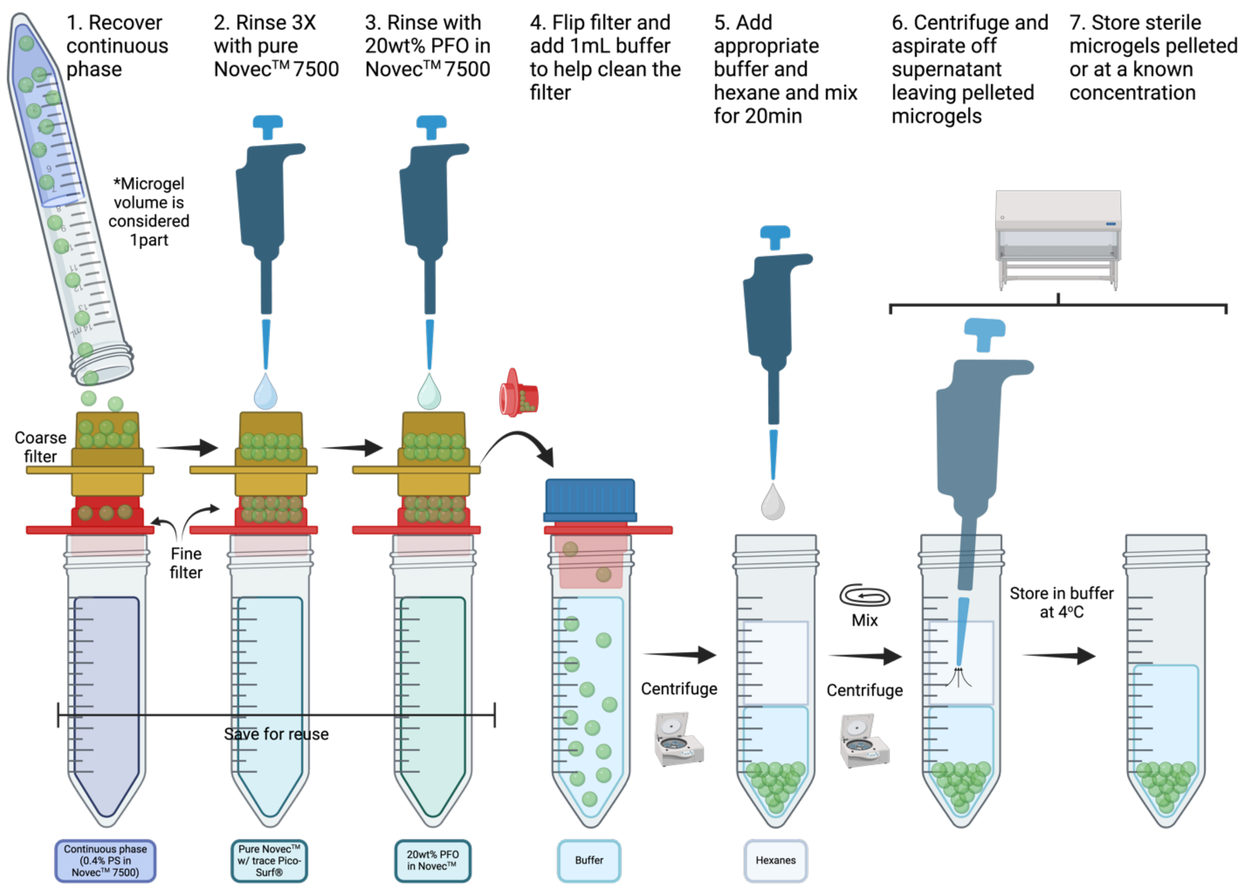


***Figure 2.*** *Overview of size-exclusion microgel purification*

#### 1. Preparation

a. Gather pluriStrainers (filters) applicable to your work.

b. Generate a 20wt% perfluorooctanol (PFO) in Novec™ 7500 solution. This will serve as the emulsion breaker. Cover in foil for storage.

c. Gather at least 4 x 50mL conical tubes.

   i. The first is for 0.4wt% Pico-Surf® in Novec™ 7500 recovery. Users may recover and reuse this solution as needed by syringe filtering through a 0.45µm syringe filter.

ii. The second is for trace Pico-Surf® in Novec™ 7500 recovery. Users may recover and reuse this solution as needed by syringe filtering through a 0.45µm syringe filter.

iii. The third is for 20wt% PFO in Novec™ 7500 recovery. Users may recover and reuse this solution as needed by syringe filtering through a 0.45µm syringe filter.

iv. The fourth is for cleaning and storage of microgels of a certain size.

v. Additional conical tubes will be needed for any other filter sizes. Aliquoting the sterile gel may be done using positive displacement pipettes and tips.

d. Gather materials in a fume hood to complete microgel purification.

#### 2. Method 1 – Size Exclusion Processing

a. For large amounts of microgels, consider parallelizing the protocol.

b. Firmly connect the pluriStrainers in order from coarsest (top) to finest (bottom). It is best to use the strainers with the taller reservoir facing up in this stage (see figure 2).

c. Place the pluriStrainers on top of the first 50mL conical tube.

d. Pour microgel solution into the top strainer to recover 0.4wt% Pico-Surf® in Novec™ 7500.

   i. Users may have to wait for solutions to pass through filters before pouring more.

   ii. To limit evaporation, place caps on the top pluriStrainer and conical tubes.

   iii. Microgels still adhered to the initial conical tube will be recovered in step 1F.

e. Once most of the liquid passes through the strainers (it will stop dripping), move the pluriStrainers containing microgels to the second 50mL conical tube (no flipping, just translating).

f. Pipette 1 mL of fresh/pure Novec™ 7500 into the original collection container and pour the solution into the top pluriStrainer. Repeat at least 3X or until sufficient amount of microgels are recovered.

   i. This rinsing step removes excess Pico-Surf® surfactant from the microgels so wait for Novec to clear the strainers before adding more.

g. Move the pluriStrainers containing microgels to the second 50mL conical tube (no flipping, just translating).

h. Once the liquid passes through the strainers (it will stop dripping), move the pluriStrainers containing microgels to the third 50mL conical tube (no flipping, just translating).

i. Slowly pipette 1mL of 20wt% PFO in Novec™ 7500 into the top pluriStrainer ensuring all the microgels come in contact with the solution. Repeat once more.

j. Once the liquid passes through the strainers (it will stop dripping), flip (one by one) the pluriStrainers containing sorted microgels to the fourth ($5^{th}$, $6^{th}$, etc.) 50mL conical tube.

   i. Keep these strainers flipped over inside new conical tubes.

k. Slowly pipette 1mL of your preferred buffer onto the new conical tube/strainer assembly and tape the conical tube cap to the top of the assembly.

l. Centrifuge this assembly at 1500rcf for 3min to remove gel from the filter.
    i. Once done, pluriStrainers may be cleaned using DI water and isopropanol.

#### 3. Hydrating and washing the microgels

a. Based on the expected volumetric swelling factor of the microgel, add the appropriate amount of buffer in 1.5X excess.
b. In the fume hood, carefully add equal parts or 15mL of Hexanes (whichever is more).
    i. Hexane lowers the density of Novec 7500, resulting in it being on the top layer where it will be aspirated.
c. Screw on the cap to the falcon tube prior to vortexing.
d. Vortex 3X for 5sec each. ***Hexanes can leak from the conical tube/cap threading so be sure to hold by the cap.
e. Place the tube on a rotating mixer so that the conical tube gets inverted during mixing but not long enough to leak through the tube/cap interface. Mix for 20min.
f. This solution is now considered “sterile”. To keep microgels sterile, complete the rest of the protocol in a biosafety cabinet (BSC) and follow sterile protocols.
g. Centrifuge at 5000rcf for 30sec. Open in BSC. Aspirate all the hexane supernatant and dispose properly.
    i. Use centrifuge vessel with lid to move into the BSC. <u>CAUTION:</u> Hexane vapors may be generated during centrifugation.
h. Centrifuge at 5000rcf for 2min. Aspirate any residual hexane supernatant and dispose properly.
    i. Aspirating a little buffer is ok since most gels are not localized at the top of the buffer/microgel phase.
    ii. Use centrifuge vessel with lid to move into the BSC. <u>CAUTION:</u> Hexane vapors may be generated during centrifugation.
i. Centrifuge at 6000rcf for 5min (depending on density difference, centrifugation time/speed may need to be increased). Aspirate excess buffer leaving concentrated microgels.
    i. This stage generates a microgel pellet within the buffer. Do not aspirate the pellet.
j. Control microgel concentration by adding known amount of buffer.
k. Aliquot with positive displacement pipette to preserve sterility.

Created 02/09/2023
Author: Durante Pioche-Lee
Lesher-Pérez Lab

### Procedure – Method 2 (No size exclusion)

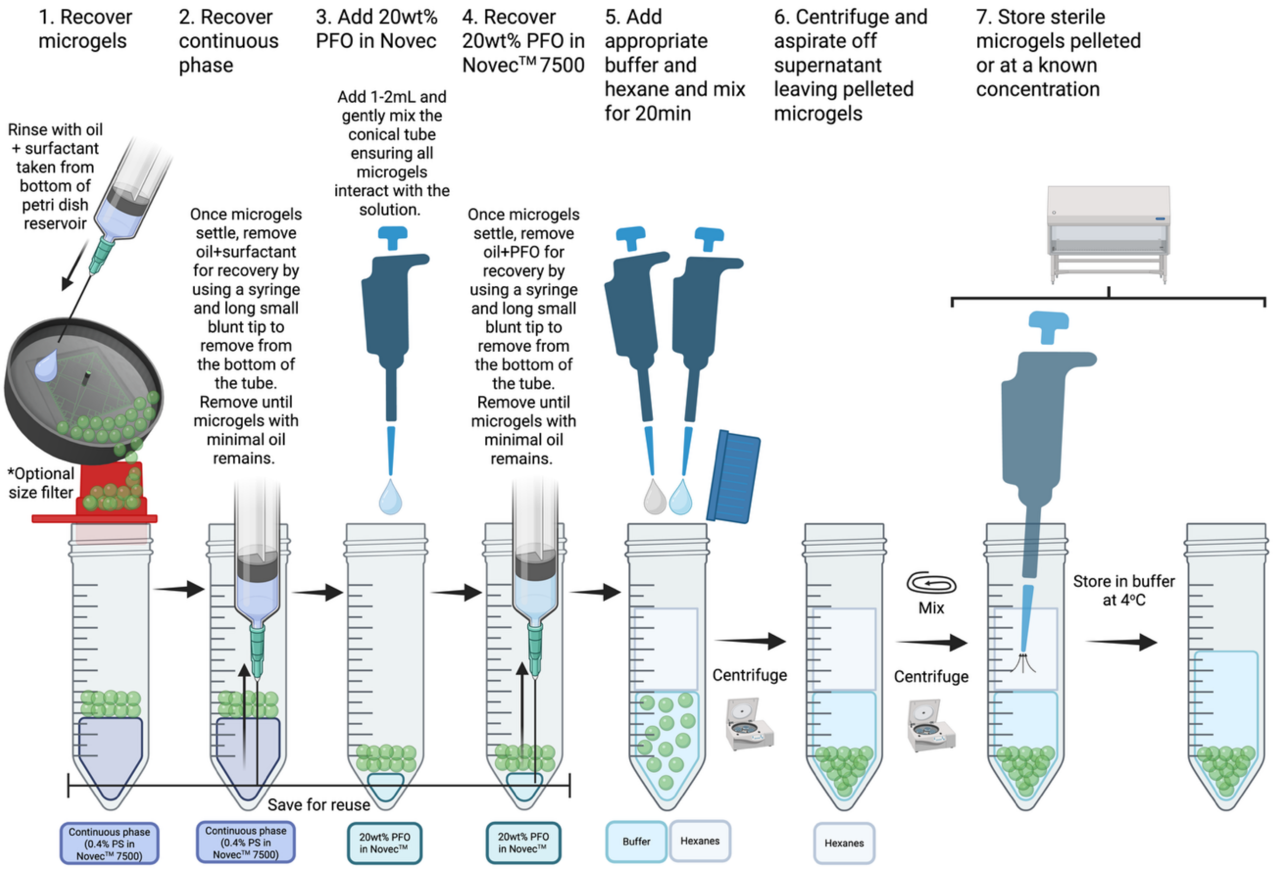


#### 1. Preparation

a. Assemble a syringe and 1.5” 22G blunt tip with 1.5”

b. Generate a 20wt% perfluorooctanol (PFO) in Novec™ 7500 solution. This will serve as the emulsion breaker. Cover in foil for storage.

c. Gather at least 3 x 50mL conical tubes.

  i. The first is for 0.4wt% Pico-Surf® in Novec™ 7500 recovery. Users may recover and reuse this solution as needed by syringe filtering through a 0.45µm syringe filter.

  ii. The second is for trace Pico-Surf® in Novec™ 7500 recovery. Users may recover and reuse this solution as needed by syringe filtering through a 0.45µm syringe filter.

  iii. The third is for 20wt% PFO in Novec™ 7500 recovery. Users may recover and reuse this solution as needed by syringe filtering through a 0.45µm syringe filter.

d. Gather materials in a fume hood to complete microgel purification.

#### 2. Method 2 – Removing non-microgel solutions

a. Centrifuge microgel container at 1500rcf for 1min. Centrifuge until there is a clear phase separation
b. For large amounts of microgels, consider using a 22G blunt tip with a longer cannula.
    i. The thin cannula on the blunt tip is used to limit microgel adhesion area when drawing from the bottom of the conical tube.
c. Place the cannula through the microgel phase and draw out the 0.4wt% Pico-Surf® in Novec™ 7500 from the bottom of the conical tube. Pay attention to not draw any microgels as this will reduce yield.
d. Wipe the blunt tip on the side of the microgel container before placing the solution into a new conical tube for 0.4wt% Pico-Surf® in Novec™ recovery.
e. Pipette 1 microgel volume of fresh/pure Novec™ 7500 into the original collection container. Centrifuge at 1500rcf for 1min. Remove the solution and place into a new conical tube for trace Pico-Surf® in Novec™ 7500 recovery. Repeat at least 1 more time.
f. After removal of the trace Pico-Surf® in Novec™ 7500. Break the emulsion by pipetting 1/4 the microgel volume of 20wt% PFO in Novec™ 7500 onto the microgels. Add 1mL of buffer and gently mix for 1min.
g. Centrifuge at 1500rcf for 1min. Remove the solution and place into a new conical tube for 20wt% PFO in Novec™ 7500 recovery.

#### 3. Hydrating and washing the microgels

a. Based on the expected volumetric swelling factor of the microgel, add the appropriate amount of buffer in 1.5X excess.
b. In the fume hood, carefully add equal parts or 15mL of Hexanes (whichever is more).
    i. Hexane lowers the density of Novec 7500, resulting in it being on the top layer where it will be aspirated.
c. Screw on the cap to the falcon tube prior to vortexing.
d. Vortex 3X for 5sec each. ***Hexanes can leak from the conical tube/cap threading so be sure to hold by the cap.
e. Place the tube on a rotating mixer so that the conical tube gets inverted during mixing but not long enough to leak through the tube/cap interface. Mix for 20min.
f. This solution is now considered "sterile". To keep microgels sterile, complete the rest of the protocol in a biosafety cabinet (BSC) and follow sterile protocols.
g. Centrifuge at 5000rcf for 30sec. Open in BSC. Aspirate all the hexane supernatant and dispose properly.
    i. Use centrifuge vessel with lid to move into the BSC. <u>CAUTION:</u> Hexane vapors may be generated during centrifugation.
h. Centrifuge at 5000rcf for 2min. Aspirate any residual hexane supernatant and dispose properly.

i. Aspirating a little buffer is ok since most gels are not localized at the top of the buffer/microgel phase.

ii. Use centrifuge vessel with lid to move into the BSC. CAUTION: Hexane vapors may be generated during centrifugation.

i. Centrifuge at 6000rcf for 5min (depending on density difference, centrifugation time/speed may need to be increased). Aspirate excess buffer leaving concentrated microgels.

   i. This stage generates a microgel pellet within buffer. Do not aspirate the pellet.

j. Control microgel concentration by adding known amount of buffer.

k. Aliquot with positive displacement pipette to preserve sterility.
